# A multi-symptomatic model of heroin use disorder in rats reveals distinct behavioral profiles and neuronal correlates of heroin vulnerability versus resiliency

**DOI:** 10.1101/2024.02.22.581440

**Authors:** Brittany N. Kuhn, Nazzareno Cannella, Ayteria D. Crow, Veronica Lunerti, Arkobrato Gupta, Stephen J. Walterhouse, Carter Allen, Reda M. Chalhoub, Eric Dereschewitz, Analyse T. Roberts, Mackenzie Cockerham, Angela Beeson, Rusty W. Nall, Abraham A. Palmer, Gary Hardiman, Leah C. Solberg Woods, Dongjun Chung, Roberto Ciccocioppo, Peter W. Kalivas

**Affiliations:** Department of Neuroscience, Medical University of South Carolina, Charleston, SC, USA; School of Pharmacy, Center for Neuroscience, Pharmacology Unit, University of Camerino, Camerino, IT; The Interdisciplinary Ph.D. Program in Biostatistics, The Ohio State University, OH, USA; Department of Biomedical Informatics, The Ohio State University, Columbus, OH, USA; Pelotonia Institute for Immuno-Oncology, The James Comprehensive Cancer Center, The Ohio State university, OH, USA; Department of Internal Medicine, Wake Forest University, Winston-Salem, NC, USA; Department of Psychology, Jacksonville State University, Jacksonville, AL, USA; Institute for Genomic Medicine, University of California San Diego, La Jolla, CA, USA; School of Biological Sciences, Queen’s University Belfast, Belfast, Northern Ireland, UK

## Abstract

**Objective:** The behavioral and diagnostic heterogeneity within human opioid use disorder (OUD) diagnosis is not readily captured in current animal models, limiting translational relevance of the mechanistic research that is conducted in experimental animals. We hypothesize that a non-linear clustering of OUD-like behavioral traits will capture population heterogeneity and yield subpopulations of OUD vulnerable rats with distinct behavioral and neurocircuit profiles.

**Methods:** Over 900 male and female heterogeneous stock rats, a line capturing genetic and behavioral heterogeneity present in humans, were assessed for several measures of heroin use and rewarded and non-rewarded seeking behaviors. Using a non-linear stochastic block model clustering analysis, rats were assigned to OUD vulnerable, intermediate and resilient clusters. Additional behavioral tests and circuit analyses using c-fos protein activation were conducted on the vulnerable and resilient subpopulations.

**Results:** OUD vulnerable rats exhibited greater heroin taking and seeking behaviors relative to those in the intermediate and resilient clusters. Akin to human OUD diagnosis, further vulnerable rat sub-clustering revealed subpopulations with different combinations of behavioral traits, including sex differences. Lastly, heroin cue-induced neuronal patterns of circuit activation differed between resilient and vulnerable phenotypes. Behavioral sex differences were recapitulated in patterns of circuitry activation, including males preferentially engaging extended amygdala stress circuitry, and females cortico-striatal drug cue-seeking circuitry.

**Conclusion:** Using a non-linear clustering approach in rats, we captured behavioral diagnostic heterogeneity reflective of human OUD diagnosis. OUD vulnerability and resiliency were associated with distinct neuronal activation patterns, posing this approach as a translational tool in assessing neurobiological mechanisms underpinning OUD.

## Introduction

Opioid use disorder (OUD) inflicts worldwide social and personal harm, with over 16 million individuals affected (1) and approximately 1 million a year seeking treatment (2). Greater understanding of the neurobiological mechanisms underlying OUD is necessary to advance treatment options. A consequential impediment towards attaining this goal is the behavioral heterogeneity inherent in a diagnosis of OUD based on DSM-V criteria. Different behavioral traits, or diagnostic criteria, interact and combine in divergent ways to form an OUD diagnosis, resulting in individuals having the same diagnosis with distinct symptom profiles. Animal models necessary for mechanistic neurobiological experimentation fail to recapitulate the multi-dimensional diagnosis of OUD and largely fail to capitalize the preclinical biological discoveries into useful treatments. We hypothesized that a non-linear clustering protocol applied to multiple behaviors commonly used in mechanistic animal studies of substance use disorder (SUD) would better recapitulate the multidimensional diagnosis of SUD and behavioral diversity conferring vulnerability. We further hypothesized this approach would yield heroin-conditioned cue circuit activation differences between OUD vulnerable and resilient subpopulations.

Typically preclinical studies use individual behavioral traits or summated linear relationships between a few traits to predict and explore underpinning neurobiological causes of SUD (3, 4). These models either do not capture a multi-symptomatic diagnosis, or when doing so rely on linear interactions that do not capture the complex multidimensional interactions between symptoms that creates subpopulations of SUDs patients. Here we describe a rat model and analysis workflow that clusters behaviors using non-linear modeling (5) on data from over 900 male and female outbred heterogeneous stock rats, a line that emulates the complex within species genetic and behavioral heterogeneity found in outbred mammalian populations, including humans (6). Rats were clustered as OUD vulnerable, resilient or intermediate. We further assessed variability within the OUD vulnerable subpopulation and identified distinct sub-clusters that, akin to human OUD, varied in combinations of traits conferring vulnerability. Also, by correlating Fos protein synthesis (indicator of neuronal activity) stimulated by heroin-conditioned cues across different brain nuclei we identified distinct patterns of connectivity between vulnerable and resilient rats.

## Methods

The Medical University of South Carolina Institutional Animal Care and Use Committee, and the Italian Ministry of Health, approved of all experimental procedures. All procedures complied with the National Institute of Health Guide for the Care and Use of Laboratory Animals and the Assessment and Accreditation of Laboratory Animals Care, as well as the European Community Council Directive for Care and Use of Laboratory Animals.

### Subjects

Heterogeneous stock rats (NMcwiWFsm:HS) were bred and shipped from Wake Forest University (WFU, USA) to either Medical University of South Carolina (MUSC, USA) or the University of Camerino (UCAM, Italy). Litters were equally represented at both sites in an effort to create genetically comparable testing cohorts. Final analyses were comprised of 917 rats (male, n=465 (MUSC, n=271; UCAM, n=194); females, n= 452 (MUSC, n=258; UCAM, n=194)). Yoked saline control rats underwent identical testing procedures as their counterparts.

### Drugs

Heroin hydrochloride supplied by the National Institute on Drug Abuse (Bethesda, MD) was dissolved in 0.9% sterile saline.

### Behavioral testing

The behavioral testing procedure is shown in Figure 1a. Rats underwent long-access heroin self-administration to quantify consumption and escalation of use. Extinction assessed perseverance of seeking in the absence of reward. Estimates of clinical laboratory-induced craving included both heroin-prime and cued reinstatement. Motivation for reward was assessed using progressive ratio test. Additional behavioral traits associated with SUD were also examined prior to and following the heroin self-administration protocol, including the elevated plus maze (EPM) and open field test (OFT) to assess stress- and anxiety-like behaviors and analgesic threshold was evaluated using the tail-flick (TF) test. TF consisted of two phases: baseline (1 mg/kg saline injection, s.c.) and test (0.75 mg/kg heroin or saline given to control rats). Additionally, a forced swim task (FST) was administered at WFU prior to shipment to assess stress-coping strategy.

**Figure 1.**
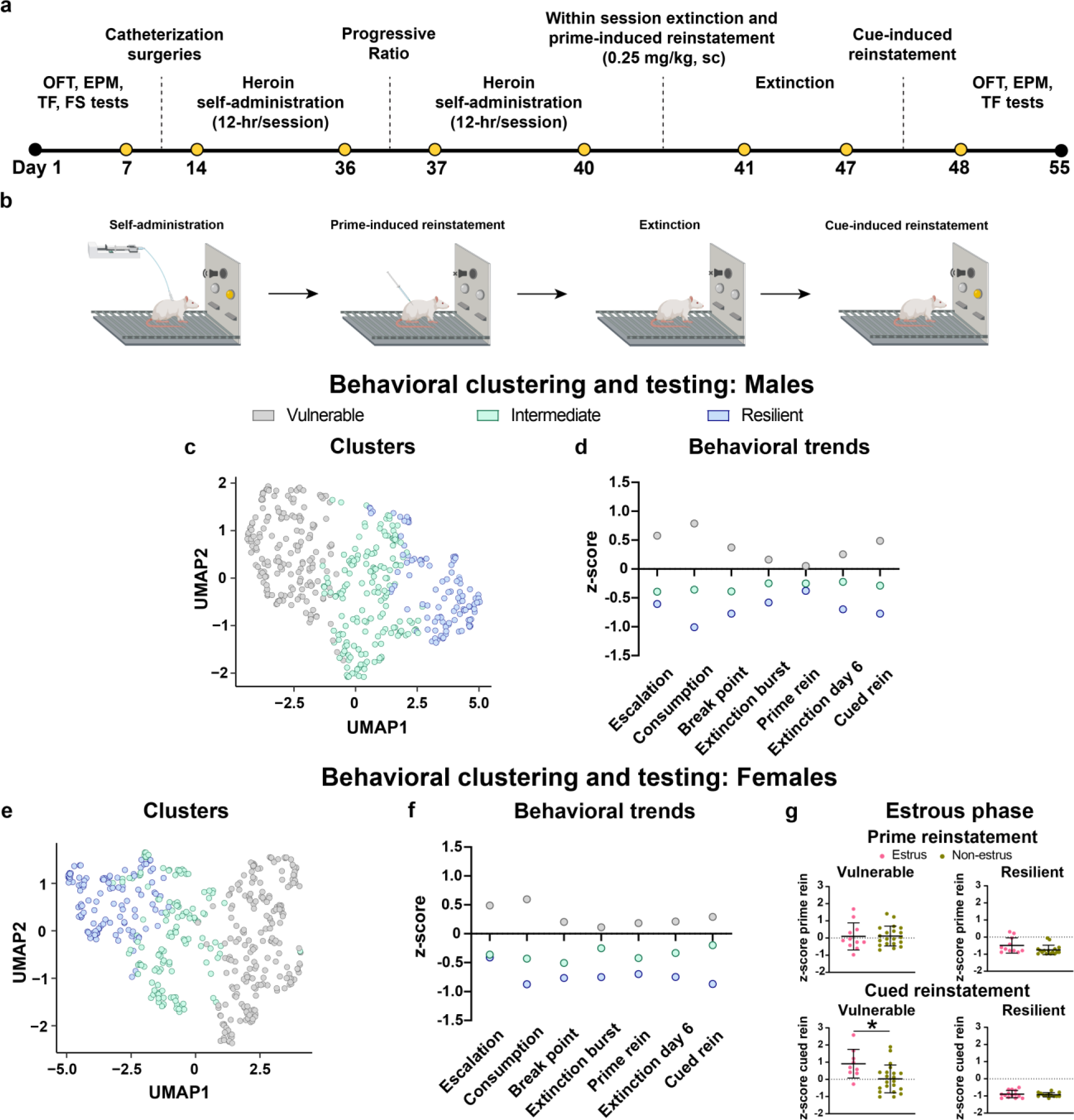
Experimental timeline and animal OUD phenotype behavioral data. **a)** Rats underwent behavioral testing (OFT: open field test; EPM: elevated-plus maze; TF: tail flick test; FS: forced swim test) prior to jugular catheterization surgery, and again at the end of heroin experience (excluding FS test). Heroin self-administration then commenced for 3 weeks (12-hr/session, 4 days a week), followed by a progressive ratio test and 3 additional days of self-administration training. A within-session extinction-prime reinstatement test then occurred, followed by extinction training and a test for cued reinstatement. **b)** Illustrations of behavioral paradigm and testing chamber configuration (created with BioRender.com). Active lever presses during self-administration training resulted in tone/cue-light presentation for 5-seconds and a heroin infusion. Active lever presses had no consequence during the within-session extinction-prime test or during extinction training. During cued reinstatement, active lever presses resulted in tone/cue-light presentation, but no heroin infusion. **c)** UMAP representation of male rats separated into distinct OUD vulnerability clusters using stochastic block model network-based clustering analysis. **d)** Median value for heroin taking, extinction and seeking behavioral measures in male rats. **e)** UMAP presentation of the distinct cluster formation of OUD vulnerability in female rats following stochastic-block model network-based clustering analysis. **f)** Median value for heroin taking, extinction and seeking behaviors between clusters in female rats. **g)** Mean ± SD for behavior during the reinstatement tests in female rats. Rats (vulnerable, n=31; resilient, n=27) in the estrus phase of the estrous cycle exhibited potentiated responding during the cued reinstatement test relative to non-estrus rats (t(28)= 2.72, p=0.01). Estrous phase cycle did not affect behavior in vulnerable rats during heroin-primed reinstatement (t(29)= 0.08, p=0.94), or in resilient rats (heroin-primed reinstatement: t(25)= 1.88, p=0.07; cued reinstatement: t(25)= 0.57, p=0.57). *p<0.05 (Males: **c-d**, Vulnerable, n= 182; Intermediate, n=168; Resilient, n=115; Females **e-f**: Vulnerable, n= 204; Intermediate, n=132; Resilient, n=116).

### Heroin self-administration, extinction and reinstatement procedures

Rats were outfitted with an indwelling jugular catheter prior to training. During self-administration presses on the active lever resulted in an infusion of heroin (20 µg/kg/100 µl infusion over 3 seconds) and presentation of a tone and light cue (Fig. 1b). Saline-control animals received a non-contingent infusion of saline (100 µl) every 20 min, accompanied with tone/cue-light presentation. There were four 12-h sessions/week for a total of 12 training sessions. A progressive ratio test occurred on day 36 (Fig. 1a), then self-administration was re-established. A 6-h extinction/prime session occurred (Fig. 1b) and rats were administered a heroin priming injection (0.25 mg/kg, s.c.; saline to control rats) after 4-h of extinction. Daily extinction training sessions then commenced prior to a test for cue-induced reinstatement (Fig. 1b). Estrous cycle phase was identified in a subset of MUSC rats (n=36) following the tests for heroin-prime and cued reinstatement.

### Additional behavioral testing

#### Punished heroin-taking behavior

We hypothesized that vulnerable rats would continue seeking heroin more than the resilient rats in the presence of an adverse stimuli. Three weeks following all testing, a subset of MUSC rats (males, n=15; females, n=14) re-established heroin self-administration training for 3 days prior to a test session under self-administration conditions where there was a 50% chance of foot shock delivery (0.40 mA, 0.5 seconds) with each infusion earned.

#### Ultrasonic vocalizations (USVs)

We examined the potential involvement that different withdrawal-induced affective states were associated with vulnerable and resilient rats. Four hours prior to extinction on day 41 (Fig. 1a), a subset of MUSC rats (males, n=22; Females, n=21; Saline, n= 8) were assessed for acute heroin withdrawal-induced USVs, a preclinical test for assessing affective state (7, 8).

#### Fos protein quantification

A subset of MUSC vulnerable and resilient rats (Females, n=13; Males, n=18; Saline, n= 11) underwent 9 additional extinction training sessions and a second cued reinstatement test ∼3 weeks after testing was completed. Fos protein expression in regions of interest were manually quantified.

### Statistical analyses

To capture behaviors across OUD phases (heroin use, extinction and seeking), seven behaviors were selected for analyses as the dependent variables: Heroin use included escalation of intake (µg/kg; average consumption days 1-3 subtracted from days 10-12) and total consumption (µg/kg) across the first 12 training sessions; Extinction behaviors include active lever presses made during the first 2 hours of the extinction-prime test (extinction burst) and during the last day of extinction training (extinction day 6); Seeking measures include break point (maximum active lever presses expended to receive an infusion) from the progressive ratio test, active lever presses during heroin-prime reinstatement and cued reinstatement.

Normality was evaluated using a Kolmogorov-Smirnov test. Raw data were assessed for group differences using a 2-way ANOVA with sex and site as the independent variables and Bonferroni post-hoc test, with several differences found (Table S1). Accordingly, data were z-score transformed within site and assessed for sex (independent variable) differences using the Mann-Whitney test (Figure S1). Sex differences were present for every behavior, and data were standardized within sex and site prior to being recombined for clustering analyses. Sexes were then analyzed as independent groups for all subsequent analyses.

We previously described a nonlinear network-based clustering analysis (9) and provided an R package ‘mlsbm’ (5). A rat-rat similarity network was constructed from the standardized data to assess behavioral similarities between rats, and then a Bayesian stochastic block model (SBM) was applied to define clusters within the data set. Cluster number was set to 3 (vulnerable, intermediate, resilient) in order to identify subpopulations of rats potentially most susceptible and least susceptible to developing OUD and in alignment with a previous study where cluster number was determined according to Bayesian Information Criterion (Figure S2) (9). Behavioral differences between clusters (independent variable) were analyzed using either an ANOVA and Bonferroni post-hoc test or Kruskal-Wallis test and Dunn’s post-hoc test.

Repeated-measures ANOVAs (cluster and session as independent variables) and Bonferroni post-hoc test were used to assess behavior during heroin-reacquisition. Unpaired Student’s-t or Mann-Whitney tests were used for punishment test, USV, and Fos protein analyses (cluster as independent variable for all) and estrous phase (phase as independent variable). To account for experimental training conditions shared across all animals that could affect Fos protein expression (e.g., exploratory behavior, testing context, etc.), Fos was standardized to saline control rats prior to analyses. Spearman correlation coefficient was used for behavioral correlations with Bonferroni test to correct for multiple comparisons and Fos protein correlations which employed a false discovery rate of q=0.01 to correct for multiple comparisons. Differences in group composition were evaluated with a Chi-squared test. Sub-clustering analyses were performed using an agglomerative hierarchical clustering strategy. The Euclidian distance between subjects with a threshold of 0.7*max(linkage) was used to create sub-clusters and a minimum of 15 rats/sub-cluster was necessary to be included in analyses. Multiple comparisons across different analyses were not corrected for. Unless stated otherwise, analyses were performed using GraphPad Prism version 9.5.1 (San Diego, CA), with p<0.05 for statistical significance.

## Results

### Raw data

Substantial site and sex differences existed in the raw data for the OUD-like traits and the behaviors quantified prior to and following heroin experience (Table S1; Figure S3-7). Female rats exhibited an overall more OUD vulnerable phenotype than male rats (Figure S5) (10). Despite standardization of experimental procedures across sites, MUSC rats showed augmented levels of heroin-related behaviors compared to UCAM rats (Figure S5). The source of these site differences presumably results from variables we did not control, such as shipping, aspects of animal housing protocols and handling, etc. (see Supplemental Methods for detailed description). Because these uncontrolled environmental factors were not quantified and the study was designed to examine environmental factors that were experimenter controlled and more related to opioid use and relapse, we z-scored the data within site and sex for further analysis (Figure S8-9). However, in future studies it would be of interest to determine which environmental differences between sites may have contributed to behavioral differences. Following standardizations, linear regression revealed ubiquitous co-variance across variables for both sites, necessitating an alternative analysis strategy to better define OUD vulnerability (Figure S10).

### OUD-like trait clustering analysis

SBM clustering was used to model the multidimensional nature of the diagnostic criteria for OUD. Three statistically distinct (Figure S2) clusters were established as vulnerable, resilient and intermediate (Male, Fig. 1c; Female, Fig. 1e), that were distributed equivalently between sexes (Figure S11). The median value of behavioral traits was higher in the vulnerable compared with resilient rats in both sexes, although the distribution of individuals showed substantial overlap between the clusters (Male, Fig. 1d and Figure S8; Female, Fig. 1f and Figure S9). Vulnerable, but not resilient, rats in estrus showed potentiated cue-reactivity compared to non-estrus phase rats. Estrous phase had no effect on behavior during heroin-primed reinstatement (Fig. 1g).

### Non-OUD traits

Stress-coping strategy or anxiety-like behavior in basal conditions were not associated with OUD vulnerability in either sex, as measured by the FS test (Figure S12) and EPM (Males, Figure S13c,f; Females, Figure S14c,f), respectively. Novelty-induced locomotion was augmented in male but not female vulnerable compared to resilient rats (Figure S13a,b), consistent with our previous finding (10) that exploratory behavior can predict OUD vulnerability in male (r^2^=0.18, p<0.001), but not female rats (r^2^=0.09, p=0.08; Figure S14a,b). Analgesic threshold was not associated with OUD vulnerability in either sex (Males, Figure S15; Females, Figure S16). However, following heroin use, female vulnerable rats had a lower analgesic threshold compared to resilient rats (Figure S16e), implying heroin-induced alterations in pain processing.

### Additional trait testing after completing the protocol in Figure 1a

#### Punishment training

Following ∼3 weeks of forced abstinence vulnerable and resilient rats readily reacquired heroin self-administration, though vulnerable animals to a greater extent (Males: Fig. 2a, Figure S17a; Females: Fig. 2d, Figure S17c). Furthermore, both female subpopulations exhibited potentiated heroin taking on the first day of reacquisition relative to the last day of heroin self-administration training, indicative of a deprivation effect described for alcohol (11) but not previously shown for opioids (Fig. 2d). During the punished heroin-taking test, vulnerable male rats endured ∼3x the number of infusion-shock pairings, and females ∼2x, prior to attenuating heroin-taking behavior to levels of resilient rats (Males, Fig. 2b, Figure S17b; Females, Fig. 2e, Figure S17d).

**Figure 2.**
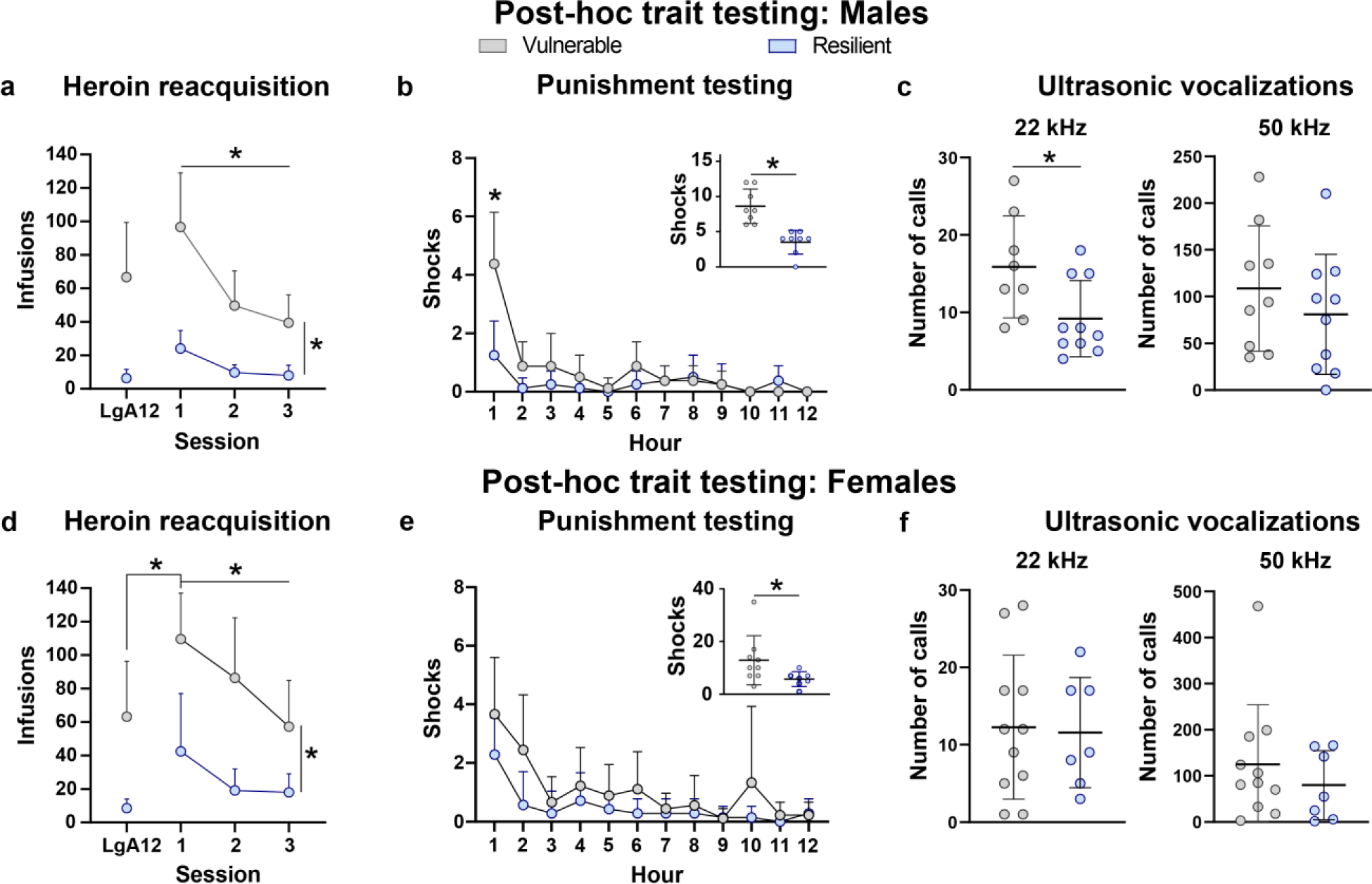
Post-hoc behavioral testing. Mean + SD **a)** for infusions earned the last day of heroin self-administration training (LgA12) and heroin reacquisition in a subset of male rats (n=8/cluster). Vulnerable male rats maintained higher levels of heroin taking during reacquisition (cluster x session interaction, F(3,42)=5.29, p=0.004; all post-hoc tests p<0.01). **b)** There was a 50% chance of foot shock delivery with each heroin infusion earned during the punishment task. Vulnerable male rats endured more shocks while maintaining heroin-taking behavior compared to resilient rats (inset, Mann-Whitney U= 0, p=0.0002), with differences centralized to the first hour of testing (cluster x hour interaction, F(11,154)=7.29, p<0.001; significant post-hoc test p=0.02). **c)** Mean ± SD for acute withdrawal-induced ultrasonic distress vocalization. Calls were higher in vulnerable male rats compared to resilient male rats (22 khz; Mann-Whitney U= 14.5, p=0.02), with no difference in appetitive state calls (50 kHz; t(17)= 0.92, p=0.37; Vulnerable, n=8; Resilient, n=10). Mean + SD **d)** for infusions earned the last day of heroin self-administration training (LgA12) and heroin reacquisition in a subset of female rats (vulnerable, n=9; resilient, n=7). A main effect of cluster, but not cluster x session interaction, was present, indicating vulnerable female rats in general maintained higher levels of heroin taking during reacquisition (cluster, F(1, 14)=36.02, p<0.0001; cluster x session interaction, F(3,42)=1.43, p=0.25), and **e)** endured more shock-heroin pairing across the test session relative to resilient rats (inset, Mann-Whitney U= 11, p=0.03). However, phenotypes only differed in cumulative shocks endured across the test session, not across the independent hours of testing (cluster x hour interaction, F(11,154)=1.56; p=0.12). **f)** Mean ± SD for acute withdrawal-induced ultrasonic vocalization calls that did not differ between female phenotypes (22 kHz, t(16)= 0.17, p=0.87; 50 kHz, Mann-Whitney U= 30, p=0.48; Vulnerable, n=11; Resilient, n=7). *p<0.05.

#### Ultrasonic vocalizations

The effect of acute heroin withdrawal on emotional state was quantified using USVs (8). USVs indicative of a positive affective state (50-kHz) did not differ between clusters in male (Fig. 2c) or female (Fig. 2f) rats. Male vulnerable rats showed potentiated USVs associated with a negative affective state (22-kHz) compared to resilient rats (Fig. 2c). Female clusters did not differ in negative affect (Fig. 2f).

### Behavioral phenotype sub-clustering

Clusters were probed for the presence of subpopulations to assess behavioral heterogeneity conferring OUD vulnerability and resiliency. Distinct sub-clusters existed only in the vulnerable phenotype for both male (Fig. 3a; n=3 sub-clusters) and female (Fig. 3b; n=4 sub-clusters) rats. For each sex, sub-clusters corresponded to augmented responding when behavior was: 1) reinforced with heroin (break point, consumption, escalation, primed reinstatement); 2) not reinforced by heroin (break point, extinction burst, extinction day 6, cued reinstatement); or 3) in both behavioral categories (Fig. 3c,d). Female rats in the non-reinforced sub-cluster exhibited further heterogeneity, with one group having potentiated responses during acute heroin withdrawal (break point, extinction burst) and the other after more protracted withdrawal (extinction day 6 and cued reinstatement; Fig. 3d). Overall, males were biased toward the heroin reinforced sub-cluster, whereas females toward the heroin non-reinforced sub-clusters (Fig. 3e). These data emulate behavioral heterogeneity in OUD diagnosis and emphasize sex differences in conferring vulnerability.

**Figure 3.**
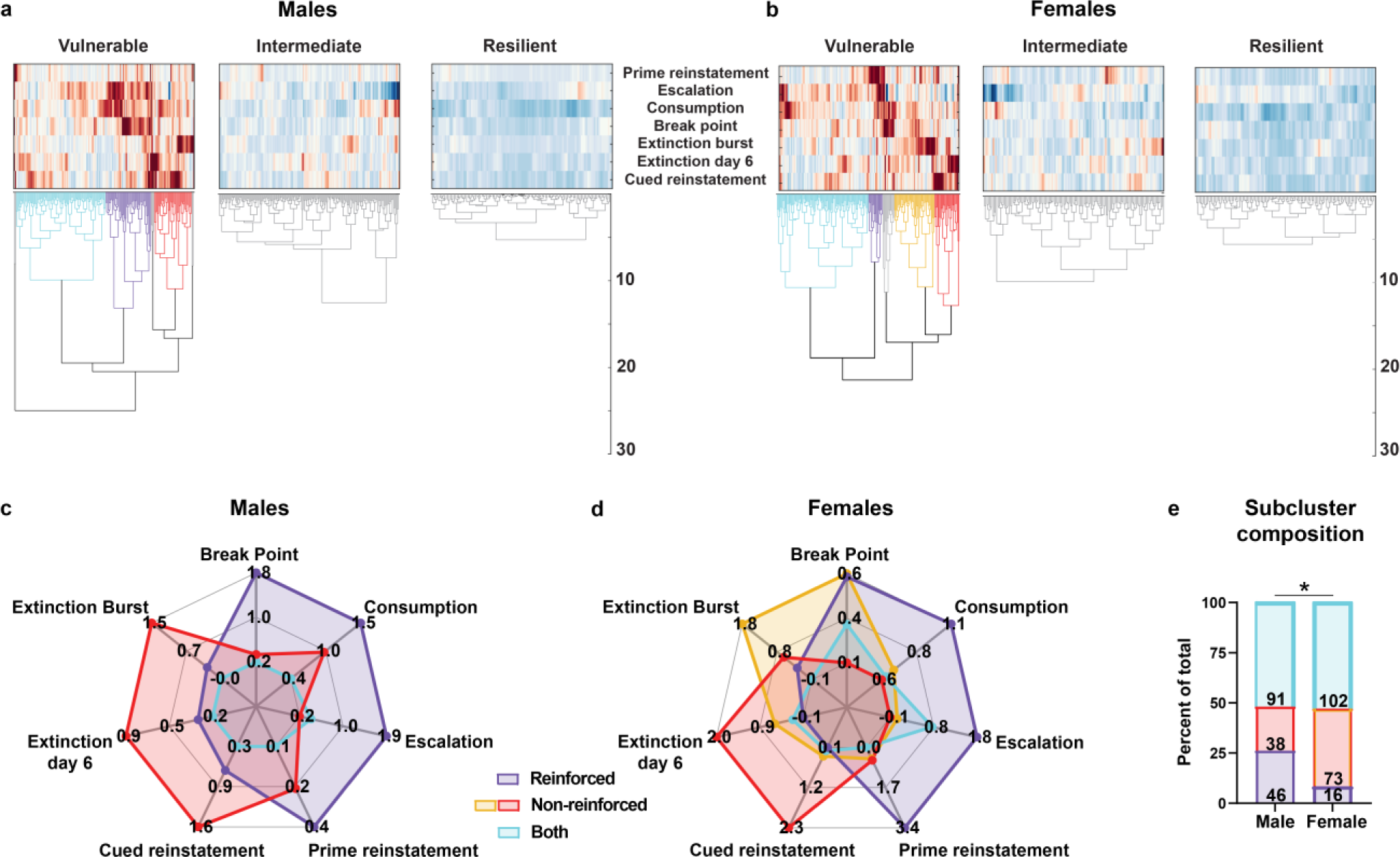
Behavioral heterogeneity present within OUD vulnerable clusters. Hierarchical analysis showed that the vulnerable group only in **a)** male and **b)** female rats was composed of distinct behavioral sub-clusters. Clusters with less than 15 animals were excluded from further analysis. Spatial representation of vulnerable sub-cluster behavioral heterogeneity presented as spider plots, where the value at each node represents the average z-score for the indicated behavior for each sub-cluster. **c-d)** Both sexes contained sub-clusters comprised of rats that showed augmented behavior when reinforced by heroin (purple), not reinforced by heroin (red/yellow) and by both circumstances (blue). **e)** Composition of sub-clusters differed between sexes, with vulnerable male rats biased toward heroin reinforced behaviors and female rats toward heroin non-reinforced behaviors (combined yellow and red sub-clusters in females; x^2^(1)=25.53, p<0.001). Number within bar is the number of rats within each sub-cluster. *p<0.05 (**a-b**, Males: vulnerable, n= 182; intermediate, n=168; resilient, n=115; Females: vulnerable, n= 204; intermediate, n=132; resilient, n=116).

### Cue-induced Fos protein expression

While group differences were not present, cue-induced Fos protein expression relative to saline rats differed between vulnerable and resilient rats in the prelimbic (PrL) and infralimbic cortex (IL), and the anterior paraventricular nucleus of the thalamus (aPVT; Figure S18-20). We were surprised to find differences in brain nuclei with opposite functions. Much of the animal and human literature show activation of PrL and IL during cued seeking (12) and of the aPVT in suppressing seeking (13, 14). We further evaluated this apparent contradiction by examining correlated Fos protein between nuclei, which revealed both shared and distinct patterns between the vulnerable and resilient phenotypes (Table S2). We subdivided ROI connectivity into subcircuits that in animal and human imaging studies (15–18) are generally associated with cue-reactivity (Fig. 4b, f), stress (Fig. 4c, g) and behavioral inhibition (Fig. 4d, h). Commensurate with vulnerable rats reinstating more that resilient rats (Mann-Whitney U= 18.50, p<0.0001; Mean ± SD: Vulnerable, 42.18 ± 25.96; Resilient, 6.07 ± 12.23; Figure S19m), vulnerable rats showed greater levels of correlated neuronal engagement (Fig. 4a,e), most notably in ROIs in the extended amygdala associated with stress responses (19) (Fig. 4c,g). It was also notable that the ventral pallidum (VP), which contains subpopulations of neurons contributing to all three subcircuits was highly interconnected within all subcircuits in vulnerable but not resilient rats. To gain insight into possible sex differences contributing toward vulnerability, vulnerable males and females were assessed separately (Table S3). Strikingly, sexes shared almost no common subcircuitry (Fig 5a, e). Female rats exhibited engagement of top-down processes, including the classic cue-reactivity pathways (20) with correlated neuronal activation between the PrL, nucleus accumbens core (NAcc) and VP (Fig. 5f). Males engaged subcortical cue-reactivity regions, specifically the NAcc and VP (Fig. 5b). Strikingly, only males showed engagement of the extended amygdala stress circuit (Fig. 5c). Since the sexes exhibited equivalent cued reinstatement (t(15)=0.27, p=0.79; Mean ± SD: Male, 44.26 ± 17.59; Female, 40.7 ± 31.40), these data suggest sexually dimorphic circuits contributing to cue reactivity.

**Figure 4.**
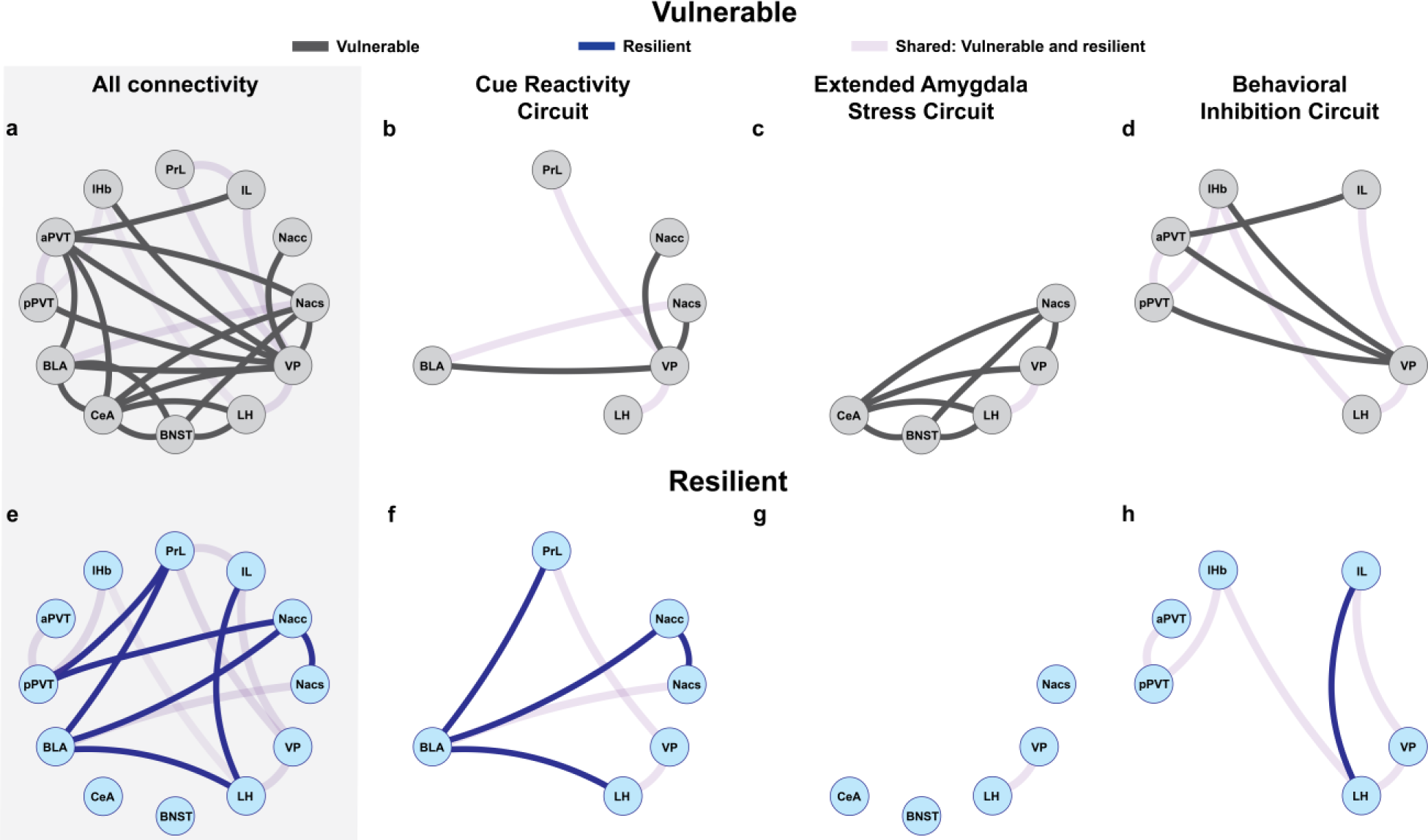
Correlated neuronal activation patterns between heroin cue-induced c-Fos protein ROIs in OUD vulnerable and resilient rats. Light purple line illustrates connectivity shared between vulnerable and resilient phenotypes; black and blue lines indicate connectivity distinct for vulnerable and resilient clusters, respectively. **a, e)** Rats in the vulnerable cluster exhibited more correlated neuronal activation compared to resilient rats. Connectivity was further broken down into three different functional circuitry categories: ROIs associated with **b, f)** cue-reactivity, **c, g)** extended amygdala stress response, and **d, h)** behavioral inhibition. Both phenotypes engaged ROIs associated with cue-reactivity, however, the vulnerable group recruited several ROIs that mediate stress response and behavioral inhibition. PrL (prelimbic cortex), IL (infralimbic cortex), NAcc (nucleus accumbens core), NAcs (nucleus accumbens shell), VP (ventral pallidum), lHb (lateral habenula), aPVT (anterior paraventricular nucleus of the thalamus), pPVT (posterior paraventricular nucleus of the thalamus), BLA (basolateral amygdala), CeA (central amygdala), LH (lateral hypothalamus), BNST (bed nucleus of the stria terminalis). Data for each ROI was standardized to saline control rats and correlations shown are p<0.05 with q=0.1 (Vulnerable, n= 17; Resilient, n=14; Saline, n=11).

**Figure 5.**
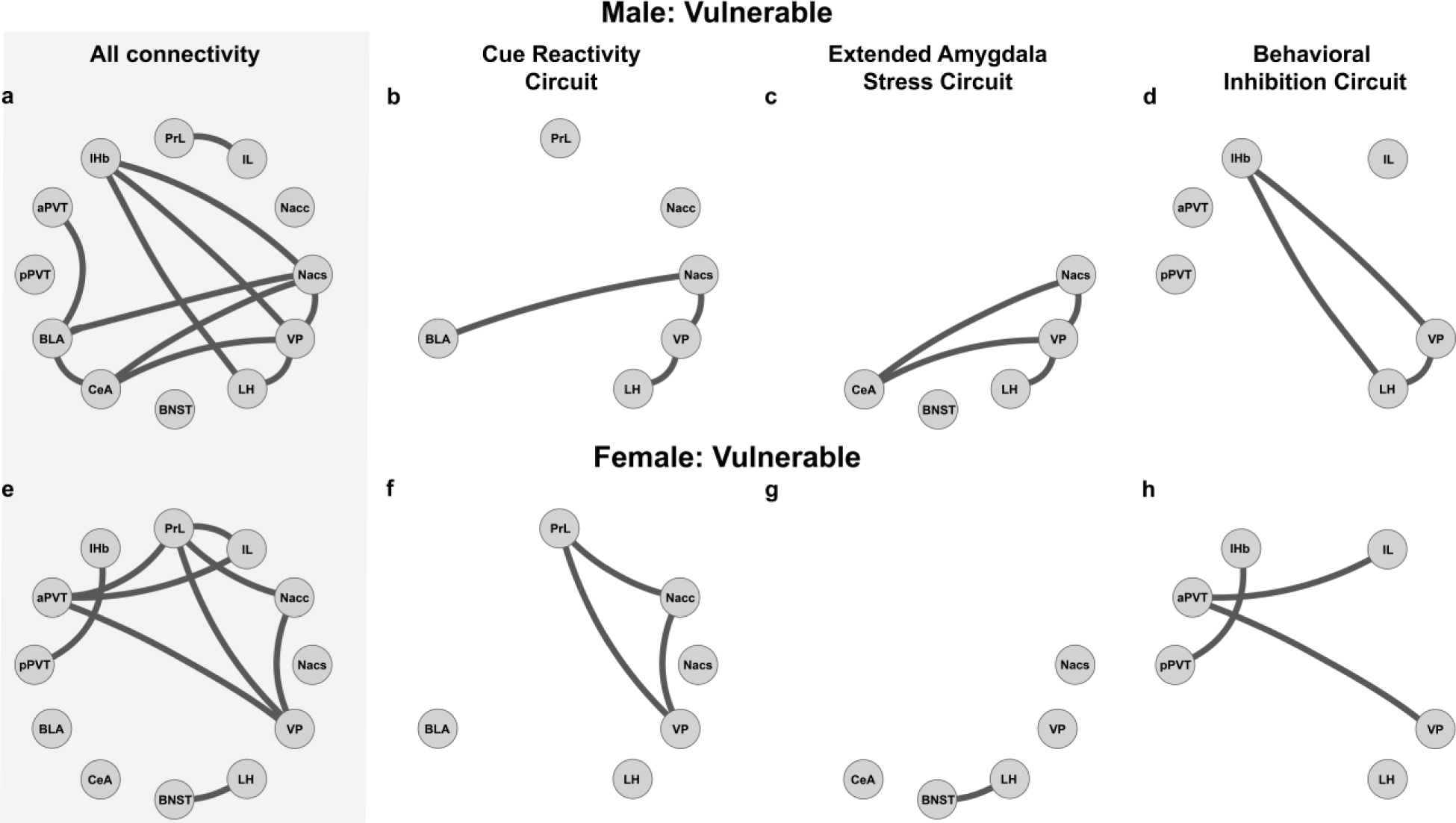
Sexual dimorphic correlated neuronal activation patterns between ROIs in male and female OUD vulnerable rats. All correlated activity patterns for **a)** male and **e)** female vulnerable rats. Connectivity patterns were further broken down into functional circuit categories: ROIs associated with **b, f)** cue-reactivity, **c, g)** extended amygdala stress response, and **d, h)** behavioral inhibition. Male rats show pronounced subcortical activity patterns, specifically engaging the extended amygdala stress circuitry. In contrast, female rats engage top-down cortical processing, with minimal engagement of stress circuitry. PrL (prelimbic cortex), IL (infralimbic cortex), NAcc (nucleus accumbens core), NAcs (nucleus accumbens shell), VP (ventral pallidum), lHb (lateral habenula), aPVT (anterior paraventricular nucleus of the thalamus), pPVT (posterior paraventricular nucleus of the thalamus), BLA (basolateral amygdala), CeA (central amygdala), LH (lateral hypothalamus), BNST (bed nucleus of the stria terminalis). Data for each ROI was standardized to saline control rats and correlations shown are p<0.05 with q=0.1 (Females, n= 10; Males, n= 7; Saline, n=11).

## Discussion

OUD has increased in all demographics over the past two decades (21), regardless of age, sex, or socio-economic status (22). To capture the diversity of OUD subpopulations and different combinations of DSM-V diagnostic symptoms we used genetically heterogeneous rats and a non-linear network-based clustering of seven rat behavioral traits. Using this approach, we succeeded in separating vulnerable from resilient rats and further sub-clustering vulnerable into subpopulations of rats possessing different combinations of OUD-like behaviors. We validated this approach by showing distinct neurocircuit activation by heroin cues between vulnerable and resilient subpopulations. We propose that this statistical workflow and behavioral findings are akin to the multi-dimensional diagnosis of OUD patients and are ideal for identifying subpopulation heterogeneity in brain mechanisms underpinning OUD vulnerability and resilience.

### OUD vulnerability clustering

The nonlinear clustering approach identified a vulnerable subpopulation of rats exhibiting greater levels of heroin use, resistance to extinction and seeking compared to rats in the intermediate and resilient clusters for both sexes. Akin to diagnostic criteria for OUD, behavioral heterogeneity was present in the traits that confer OUD-like vulnerability in both sexes, with distinct subpopulations exhibiting behavioral vulnerability in heroin reinforced drug-seeking, non-reinforced drug-seeking or a combination of both. Male vulnerable sub-clusters were biased toward heroin reinforced seeking, and females toward heroin non-reinforced seeking. Female rats in the non-reinforced subpopulation were further subdivided into a subpopulation more motivated to respond during early (i.e., progressive ratio and extinction burst) versus late extinction conditions (i.e., extinction day 6), with the latter also exhibiting high cue reactivity during cued reinstatement, suggesting differential sensitivity to contextual and discrete drug-associated cues.

### Other measures associated with vulnerability

In addition to the seven traits used for SBM clustering, hallmark features of OUD were identified in subsequent testing done in a portion of rats. Vulnerable rats re-acquired heroin self-administration and required more infusion-shock pairings to attenuate heroin-taking behavior. Male vulnerable rats exhibited more distress than their resilient counterparts during acute heroin withdrawal, with no differences observed in females. However, male rats are more prone to emit stress-induced USVs (23), suggesting alternative withdrawal assays may be more applicable for female rats. Circulating ovarian hormones affected behavioral heterogeneity in female rats. Behavioral responding is potentiated during the estrus phase of the cycle due to estradiol regulation of mesolimbic dopamine (24), and vulnerable estrus phase rats showed potentiated cued seeking relative to non-estrus phase rats. No differences were observed during prime reinstatement, suggesting estradiol involvement in modulating discrete environmental cues, but not heroin interoceptive cues.

### Correlated Fos activity patterns in OUD vulnerable versus resilient rats

Little is known about the neurobiological mechanisms mediating SUD resilience. Decline in prefrontal cortex grey matter volume (25) and engagement of top-down cortical processes within the dorsal striatum (26) confers SUD resiliency. Resilient OUD rats exhibit hypo-function of neuronal activation in medial prefrontal cortex (Figure S19), suggesting that resiliency may be mediated by prefrontal cortex neuroplasticity. Resilient rats appear to be recruiting circuitry and regions engaged in negative affective states and avoidance, such as the PrL-BLA pathway that mediates anxiety-like behavior and fear memory acquisition (27) and the IL, a region contributing toward suppressing drug seeking (28). This profile of correlated activation suggests that OUD resiliency is in part governed by circuits that inhibit behavioral responding to cued seeking.

Commensurate with greater cue-induced heroin seeking, vulnerable rats engaged more complex circuitry, similar to what is observed in human OUD individuals (16). Key features of connectivity distinctions between clusters include a majority of connectivity in vulnerable rats centered around the aPVT, a region involved in mediating anxiety and stress response (29) and suppression of reward seeking (13), the VP which exhibits cell specific regulation over drug-seeking and drug-refraining (i.e., withholding of seeking) (30), and the CeA, part of the extended amygdala that mediates stress response and stress-induced reinstatement (31). In an effort to disentangle the differences in circuit activation we isolated nuclei generally found activated in the animal and human imaging literature during a cue-reactivity test, stress responding with a focus on the extended amygdala, and behavioral inhibition (16–19). Most striking was the involvement of the extended amygdala stress circuit in vulnerable, but not resilient rats. Importantly, stress facilitates relapse in animal models and humans (32), which can account for the robust cued relapse in vulnerable, but not resilient rats. Surprisingly, the vulnerable rats also showed greater overall connectivity in behavioral inhibition circuitry, notably in the interconnectivity between the lHb, PVT and VP, all structures experimentally implicated in a cell, region or circuit-specific manner to promote or induce avoidance behaviors (14, 30, 33). It is possible that following chronic heroin self-administration in vulnerable rats, these brain nuclei have undergone adaptations rendering them less capable of inhibiting cued seeking (34), or a shift in cell-specific opposing regulation of drug seeking versus avoidance may have occurred (35).

### Sex differences

In parallel with behavioral sex differences frequently observed in this study, we found minimal overlap in vulnerability circuitry in male or female rats. Approximately 40% of female rats were in the non-heroin reinforced sub-cluster, and females heavily engaged the PrL◊NAcc◊VP series circuit that mediates cue reactivity in animals and humans (20). In contrast, male rats predominately engaged extended amygdala circuitry involved in stress responding (31). Vulnerable males also exhibited withdrawal-induced distress posing the possibility that male rats were experiencing more distress following prolonged heroin abstinence, resulting in stress circuitry contributing to cued reinstatement. Females, however, showed lower levels of distress across training. Together, these data suggest sexual dimorphism in neuroadaptations governing vulnerability in male and female rats, with male vulnerability driven more by stress circuitry and female vulnerability by cue responsivity. Importantly, the literature supports nuanced sex differences translating preclinical to human OUD studies (24), though a more focused approach is necessary in preclinical and clinical research to sufficiently disentangle SUD sex differences (36).

## Conclusions and Future Directions

Medication-assisted treatment is the current standard of OUD treatment (37). However, diagnostic heterogeneity is an important consideration for treatment, as individual symptom clusters are likely driven by distinct neurobiological mechanisms and genetics. By using a non-linear clustering model that allowed us to capture aspects of the behavioral heterogeneity and diagnostic diversity in human OUD, we identified distinct behavioral profiles and sexually dimorphic neurobiological circuitry associated with OUD resiliency versus vulnerability in rats. Further sub-clustering revealed variability in traits conferring vulnerability, with notable differences in sub-cluster composition between sexes. This study constitutes a first step in applying non-linear clustering of SUD vulnerability traits and provides a preclinical database for continued studies into the neurobiological mechanisms and genetic vulnerabilities contributing to OUD phenotypes. Additionally, neurobiological underpinnings of sex differences in human SUDs is not well characterized, and despite rates of OUD steadily increasing in females (38), females remain underrepresented in clinical OUD research (<15% of participants) (16). The marked behavioral and circuit sex differences we found highlights using non-linear clustering in animal models to discern mechanisms of sex differences in behavioral phenotypes contributing to human OUD.

## Acknowledgements

The experiments were designed by BNK, NC, AAP, GH, DC, RC and PWK, and conducted by BNK, NC, ADC, VL, SJW, ED, ATR, MC, AB and RWN. Statistical modeling was performed by CA, AG, DC, and RMC. LCSW provided all heterogeneous stock rats used in these experiments. Data was analyzed by BNK, CA, AG, DC and RMC. BNK and PWK wrote the manuscript. We would like to thank Dr. Carmela Reichel for providing the equipment necessary to perform the punished heroin taking test, and the Department of Neuroscience Behavioral Core at Medical University of South Carolina for the equipment needed to record ultrasonic vocalizations.

## Supplemental methods

### Subjects

Heterogeneous stock rats from NIH (N/NIH-HS) [1] bred at Wake Forest University (WFU) since 2017 (Rat Genome Database number 13673907) were used in the experiments. Rats were shipped from WFU (n=1200) to MUSC or UCAM at 4-6 weeks of age in batches of 40 (20 females and 20 males/site). Care was taken to ensure that each site received equivalent number of animals from each litter. Animals arrived at MUSC via van transport the same day of shipping from WFU. Rats were shipped via air travel same day from the United States to Rome, Italy, and required 48 hours of processing and van transit before arriving at UCAM. During transit, all animals had access to food and hydrogel, and were kept in a climate controlled environment. Upon arrival at both sites, rats were left undisturbed for 3 weeks in a climate-controlled vivarium (standard 12-hr light:dark cycle) prior to testing. Rats were pair-housed and had ad libitum access to standard chow (MUSC: Lab Diet 5V75; UCAM: Mucedola 4RF18) and water throughout testing. Housing chambers at MUSC contained 1204.5 cm^2^ floor space and 22,885.5 cm^3^ total volume (36.5 × 33 cm base by 19 cm tall), contained corncob bedding and were attached to a Lixit system for water delivery and ventilation. At UCAM, housing chambers were of similar dimensions at 1126.3 cm^2^ floor space and 21,398.8 cm^3^ total volume (26.5 × 42.5 cm base by 19 cm tall), contained wooden dust free bedding and were kept on a free standing rack with 2 water bottles per cage. The following animals were excluded from analyses: 107 due to death (surgery, n= 23; illness, n= 84), 29 due to technical issues in data collection, and 147 were used for control self-administration procedures (saline self-administration, n= 111; sucrose, n=36).

Rats at MUSC had to be transported via freight elevator daily from the floor containing the housing facility to the testing suite. At UCAM, the housing and testing facilities were located near-adjacent on the same floor. All testing occurred during the dark phase with the exception of the elevated-plus maze, open field test and the tail flick test (Figure 1a). These behavioral assays occurred in the following sequence starting at 07:00, with an hour between each test whereupon rats were left undisturbed in their home cage: elevated-plus maze, open field test then tail flick test. Animals were handled daily the 3 days prior to beginning the behavioral protocol. All testing procedures, including equipment used and general housing conditions, were standardized between sites.

### Behavioral testing

#### Forced swim (FS) test

Sessions lasted for 6 minutes and the last 5 minutes of each session was manually scored in 5 second intervals for measures of immobility (e.g., floating) and mobility (e.g., swimming and climbing). Testing apparatus was a container (diameter 28 cm, height 36 cm) with a water depth 27 cm and temperature of 25°C ± 2. Subsequent behavioral testing occurred 3 weeks after the rats’ arrival to the study site location, immediately preceding heroin self-administration training.

#### Elevated-plus maze (EPM)

The EPM test lasted for 5 minutes. The apparatus (San Diego Instruments; 4 arms at 110.5 cm long and 10.16 cm wide) consisted of two enclosed arms (30.48 cm high walls; “closed” arms) and two “open” arms (i.e. not enclosed). Testing occurred under dark conditions, with infrared lighting overhead of the apparatus to allow for accurate scoring. Behavior was recorded and automatically scored using ANY-maze behavioral tracking software (Stoelting, Wood Dale, IL; version 6.17) with a threshold of 85% of animal’s body within an arm to be counted in it.

#### Open field test (OFT)

Sessions lasted for 60 minutes in a Plexiglas chamber sitting within a metal frame (Omnitech Electronics, Columbus, OH; 40.64 cm L x 40.64 cm W x 30.48 cm H) housed within a dark room. Photocells within the chambers tracked lateral and vertical movements, and data was recorded and analyzed using Versamax (Omnitech Electronics, Columbus, OH; version 1.80-0142).

#### Tail flick test

Testing occurred on a platform outfitted with a motion sensor that turns the beam off once the animal removed its tail, or after 10 seconds passed (Ugo Basile S.R.L., Gemonio, Italy). Injections (i.e., saline or heroin sc) were administered 15 minutes prior to session start. The two phases were separated by 1 hour and consisted of 4 trials that were averaged to determine the mean latency to remove the tail from the sensor. To prevent tissue damage, the location of the rat’s tail over the beam was adjusted by 1 cm each trial.

### Heroin self-administration, extinction and reinstatement procedures

Prior to training, rats were outfitted with an indwelling jugular catheter and administered an analgesic (Ketorolac, 2 mg/kg, s.c.; or Meloxicam, 0.5 mg/rat, s.c.) and an antibiotic (Cefazolin, 0.2 mg/kg, s.c.; or enrofloxacin, 1 mg/kg, i.v.) post-operatively. Catheter patency was verified prior to training starting via blood collection for additional analyses (data not presented in this manuscript). If a rat showed a sudden decrease in heroin-taking behavior, or the catheter port was compromised (e.g., leaking), patency was checked using methohexital sodium diluted in sterile saline (10 mg/ml, i.v.). Rats were re-catheterized (MUSC, n= 15; UCAM, n= 30) if ataxia was not observed within 5 seconds. Following a 3-day recovery period, rats were reintroduced back into experimentation. All training occurred in standard behavioral testing chambers (Med Associates, St. Albans, VT) comprised of two levers with a light above each, and a speaker and house light on the opposite wall (Figure 1b). A fixed-ratio 1 schedule of reinforcement was used for self-administration. Once an infusion was earned a 20-second time out period was signaled by the house light turning off whereupon additional active lever presses were recorded, but did not result in a heroin infusion. Sessions terminated after 12 hours or 300 earned infusions, with four training sessions/week Monday-Friday, one randomized day off per week. A progressive ratio test was used to assess motivation to take heroin, during which the number of lever presses needed to receive an infusion exponentially increased according to the following formula: (5*e^0.2n^)-5 [2]. Testing terminated after 1 hour of no earned infusion or 12 total hours of testing. During extinction training, active lever presses were without consequence (i.e., no infusion or cue presentation). Active lever presses during cued reinstatement resulted in tone/cue-light presentation, but no infusion.

### Estrous cycle phase identification

Vaginal lumen samples were collected and processed with a hematology stain from a subset of female MUSC rats (n=36) following the tests for heroin-prime and cued reinstatement to determine estrous cycle phase at time of testing. Phase was determined via cell morphology using a Leica Thunder DM6B scope.

### Ultrasonic vocalizations

Rats were placed into a self-administration testing chamber housed within a sound attenuating cabinet in a quiet room, and ultrasonic vocalizations (USVs) were recorded for 5-minutes. Testing chambers were off (i.e., no levers or cues were present). The recording microphone was placed approximately 12.25 cm from the top of the chamber, beyond physical reach from the rat. Vocalizations were recorded using Avisoft UltraSoundGate equipment (Avisoft Bioacoustics, Glienicke/Nordbahn, Germany), and the following parameters were used: FFT was set to 256, bandwidth of 3672 Hz and resolution of 977 Hz. Recordings were sorted using Avisoft SAS Lab Pro. Both the audio recording and the spectrogram were used to identify USVs and associated frequency range, as well as to exclude ambient noise (e.g., animal locomotive behaviors, other environmental noises, etc.). USV duration had to exceed 25 ms to be included in analysis. USVs between 18-33 kHz (i.e., 22-kHz range) and 38-80 kHz (i.e., 50-kHz range) frequency range were quantified in 60-second intervals (i.e., five 60-sec bins).

### Behavioral phenotype subclusters

Sub-clustering analyses were performed using an agglomerative hierarchical clustering strategy on individual sub-clusters, using custom MATLAB scripts (not publicly available). Similarity between subjects was measured using Euclidian distance (pdist function) along the variables by which the animals were originally clustered: escalation of intake, total consumption, break point, and active lever presses during extinction burst, extinction day 6, heroin-prime reinstatement and cued reinstatement. A hierarchal clustering tree was subsequently created by the ward linkage method on the calculated distances between subjects. The tree was then divided into subsequent sub-clusters using a threshold of 0.7*max(linkage). A minimum of 15 rats per sub-cluster was used as a cut-off to be included in post-hoc analyses on the generated sub-clusters.

### Fos protein quantification

Immediately following the second reinstatement test (2-hr session), animals underwent transcardial perfusion and brains were extracted and post-fixed prior to being sliced (40 µm) for Fos protein immunohistochemistry. Brains were first washed in 0.1 M phosphate-buffered saline (PBS; 3×5 min/wash) and blocked in 1% hydrogen peroxide in PBS for 10 min prior to undergoing 4 additional PBS washes. Slices were blocked in PBS + 0.4% Triton-X + 2.5% normal donkey serum (Jackson ImmunoResearch, Product #017-000-121) for 1 hour at room temperature, then incubated overnight in rabbit anti-cFos primary antibody (1:2000 dilution; Cell Signaling, Product #5348S). Brain slices were rinsed in PBS and incubated in biotinylated donkey anti-rabbit secondary antibody (1:500 dilution; Jackson ImmunoResearch, Catalog #711-065-152) at room temperature for 1 hour and then rinsed in PBS. Incubation in ABC-Elite (1:1000 dilution; Vector Labs, Product #PK-6100) followed for 1 hour and tissue was rinsed in PBS. Last, slices were incubated for 8 minutes in diaminobenzidine tetrahydrochloride (Signma-Aldrich, Product #D5905) and 0.012% hydrogen peroxide prior to additional rinses in PBS. Slices were mounted onto slides, underwent graduated ethanol dilution washes and a Xylene wash prior to being cover slipped with Permount mounting medium.

Images of regions of interest (ROI) were acquired as a 20x brightfield tilescan using a Leica Thunder DM6B scope outfitted with a Flexacam C1 camera. ImageJ was used for manual c-Fos protein cell quantification, and ROI parameters were identical for a region across animals to standardize data. Fos protein number for each animal and ROI was standardized to the average Fos protein number in saline control rats for the respected ROI (i.e., Fos protein for ROI/average Fos protein level in saline control rats for ROI). Fos protein levels across the ROIs were correlated with one another [3-5] to assess potential neural circuitry involved in OUD vulnerability versus resiliency.

---

**Table S1.** Raw data statistics for all behavioral testing and heroin taking, extinction and seeking measures. **a)** Behavioral differences were highly prevalent between sites (MUSC vs UCAM) and sex (male vs female) both prior to (test 1) and following (test 2) heroin self-administration training. Forced swim testing only occurred prior to heroin experience. **b)** Differences were present between sites for every measure except active lever pressing during heroin-prime reinstatement. Behavior between males and females were also present, with sexes not differing to escalation of heroin taking and break point achieved during the progressive ratio test only. No interactions were present. Abbreviations: DF, degrees of freedom; F, F-test value; P, probability value.

| **a** | **Raw data: Behavioral tests** |  |  |  |  |  |  |  |  |
| --- | --- | --- | --- | --- | --- | --- | --- | --- | --- |
|  | **Site** |  |  | **Sex** |  |  | **Site \* Sex** |  |  |
|  | DF | F | P | DF | F | P | DF | F | P |
| *Tail flick baseline 1* | 1,878 | 46.84 | <0.0001 | 1,878 | 2.56 | 0.11 | 1,878 | 5.01 | **0.03** |
| *Tail flick test 1* | 1,878 | 533.3 | <0.0001 | 1,878 | 80.50 | <0.0001 | 1,878 | 20.74 | <0.0001 |
| *Tail flick difference 1* | 1,878 | 320.5 | <0.0001 | 1,878 | 58.17 | <0.0001 | 1,878 | 27.63 | <0.0001 |
| *Time travelled 1* | 1,874 | 2257 | <0.0001 | 1,874 | 2.37 | 0.12 | 1,874 | 26.40 | <0.0001 |
| *Distance travelled 1* | 1,874 | 61.76 | <0.0001 | 1,874 | 119.0 | <0.0001 | 1,874 | 56.95 | <0.0001 |
| *Percent time open arm 1* | 1,877 | 126.0 | <0.0001 | 1,877 | 83.14 | <0.0001 | 1,877 | 1.90 | 0.17 |
| *Number of climbs* | 1,843 | 0.16 | 0.69 | 1,843 | 26.66 | <0.0001 | 1,843 | 0.10 | 0.75 |
| *Time swimming* | 1,843 | 4.56 | **0.043** | 1,843 | 1.43 | 0.23 | 1,843 | 0.05 | 0.82 |
| *Time floating* | 1,843 | 3.81 | 0.05 | 1,843 | 0.69 | 0.41 | 1,843 | 0.04 | 0.84 |
| *Tail flick baseline 2* | 1,878 | 103.6 | <0.0001 | 1,878 | 5.74 | **0.02** | 1,878 | 14.36 | **0.0002** |
| *Tail flick test 2* | 1,878 | 287.4 | <0.0001 | 1,878 | 21.26 | <0.0001 | 1,878 | 9.29 | **0.002** |
| *Tail flick difference 2* | 1,878 | 136.7 | <0.0001 | 1,878 | 11.18 | **0.0009** | 1,878 | 21.02 | <0.0001 |
| *Time travelled 2* | 1,872 | 1496 | <0.0001 | 1,872 | 0.13 | 0.72 | 1,872 | 16.79 | <0.0001 |
| *Distance travelled 2* | 1,872 | 125.6 | <0.0001 | 1,872 | 47.29 | <0.0001 | 1,872 | 15.67 | <0.0001 |
| *Percent time open arm 2* | 1,877 | 4.03 | **0.04** | 1,877 | 52.68 | <0.0001 | 1,877 | 0.31 | 0.58 |

**Table S1** Raw data statistics for all behavioral testing and heroin taking, extinction and seeking measures.
| **b** | **Raw data: Heroin taking, extinction and seeking** |  |  |  |  |  |  |  |  |
| --- | --- | --- | --- | --- | --- | --- | --- | --- | --- |
|  | **Site** |  |  | **Sex** |  |  | **Site \* Sex** |  |  |
|  | DF | F | P | DF | F | P | DF | F | P |
| *Escalation* | 1,913 | 52.17 | <0.0001 | 1,913 | 2.98 | 0.08 | 1,913 | 0.46 | 0.50 |
| *Consumption* | 1,913 | 29.35 | <0.0001 | 1,913 | 66.53 | <0.0001 | 1,913 | 0.003 | 0.96 |
| *Break point* | 1,913 | 118.4 | <0.0001 | 1,913 | 1.31 | 0.25 | 1,913 | 0.11 | 0.74 |
| *Extinction burst* | 1,913 | 114.4 | <0.0001 | 1,913 | 5.46 | **0.02** | 1,913 | 0.0006 | 0.98 |
| *Prime reinstatement* | 1,913 | 0.73 | 0.39 | 1,913 | 15.69 | <0.0001 | 1,913 | 0.68 | 0.41 |
| *Extinction day 6* | 1,913 | 17.13 | <0.0001 | 1,913 | 6.09 | **0.01** | 1,913 | 1.14 | 0.29 |
| *Cued reinstatement* | 1,913 | 21.18 | <0.0001 | 1,913 | 15.89 | <0.0001 | 1,913 | 3.09 | 0.08 |

**Table S2.**
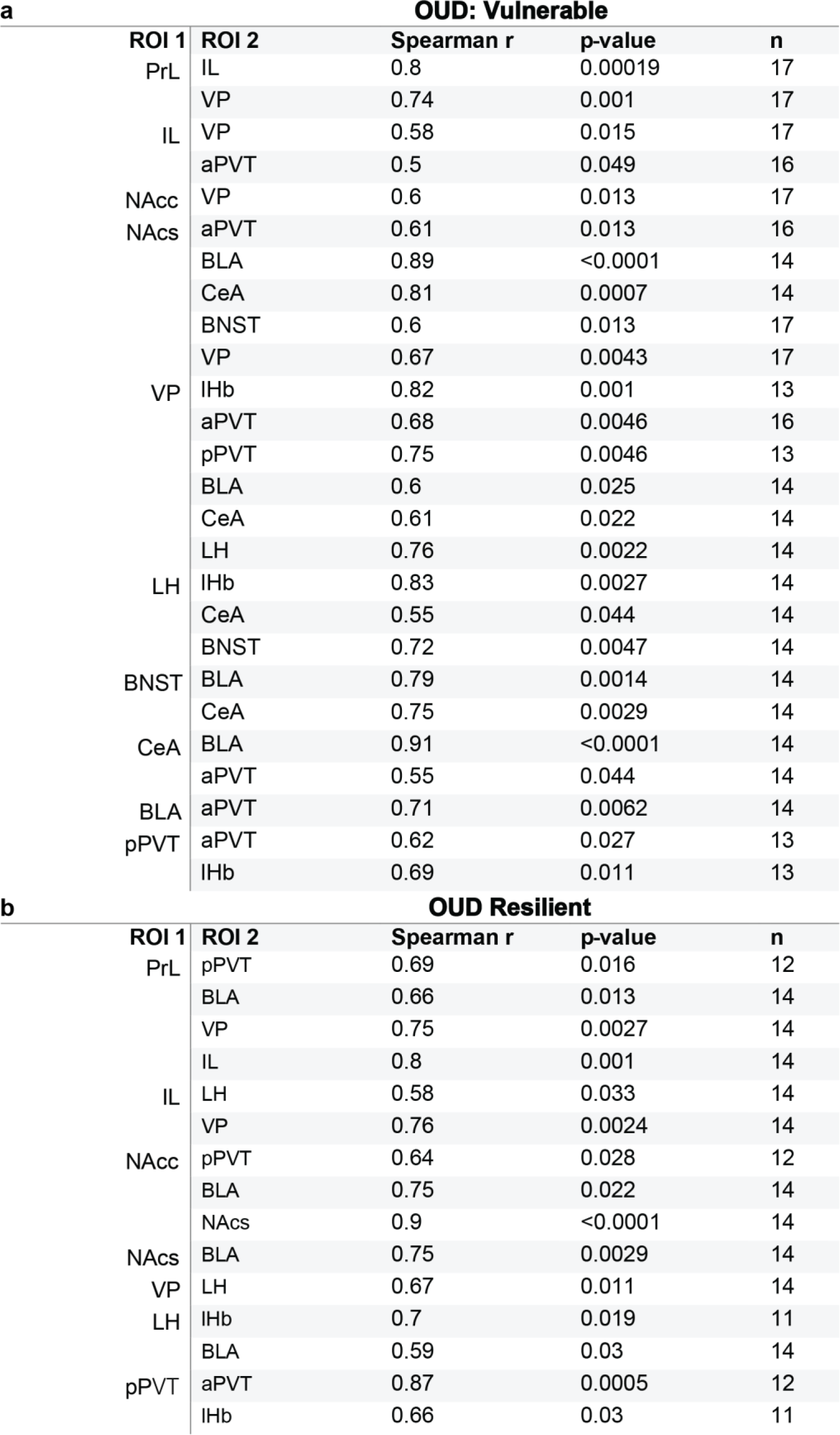
Data statistics for significant correlated neuronal activation in OUD vulnerable and resilient male and female rats. **a)** Vulnerable rats showed overall augmented levels of activation patterns relative to **b)** resilient rats. All data was standardized to saline control animals (n=11) prior to comparison. All correlations were positive, with a p<0.05 and a q=0.01 to correct for multiple comparisons. Abbreviations: ROI, region of interest; Spearman r, correlation coefficient; n, number of rats/comparison; PrL (prelimbic cortex), IL (infralimbic cortex), NAcc (nucleus accumbens core), NAcs (nucleus accumbens shell), VP (ventral pallidum), lHb (lateral habenula), aPVT (anterior paraventricular nucleus of the thalamus), pPVT (posterior paraventricular nucleus of the thalamus), BLA (basolateral amygdala), CeA (central amygdala), LH (lateral hypothalamus), BNST (bed nucleus of the stria terminalis)

**Table S3.**
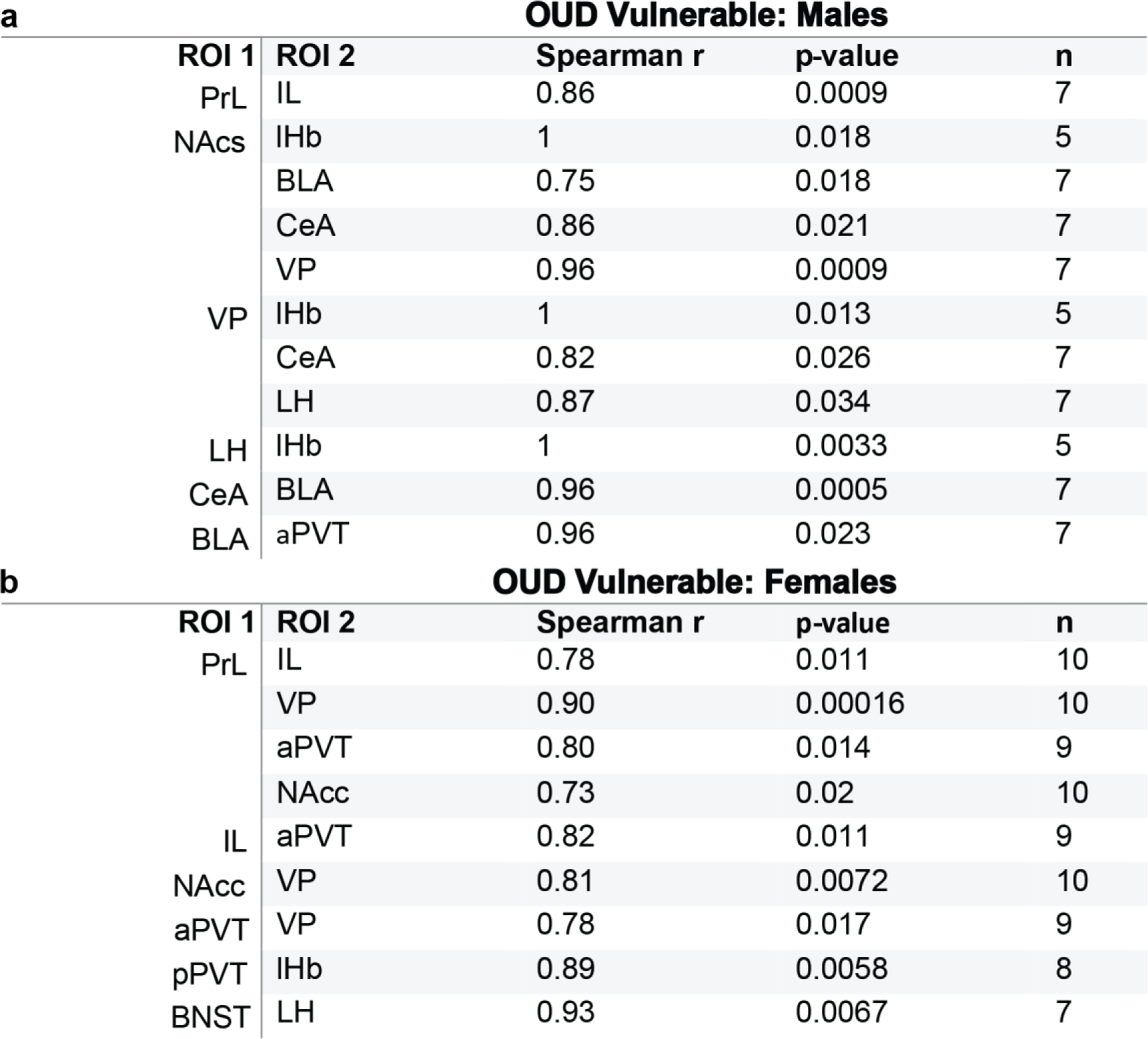
Data statistics for significant correlated neuronal activation in OUD vulnerable male and female rats. Vulnerable **a)** male and **b)** female rats showed distinct neuronal activation patterns. All data was standardized to saline control animals (n=11) prior to comparison. All correlations were positive, with a p<0.05 and a q=0.01 to correct for multiple comparisons. Abbreviations: ROI, region of interest; Spearman r, correlation coefficient; n, number of rats/comparison; PrL (prelimbic cortex), IL (infralimbic cortex), NAcs (nucleus accumbens shell), VP (ventral pallidum), lHb (lateral habenula), aPVT (anterior paraventricular nucleus of the thalamus), BLA (basolateral amygdala), CeA (central amygdala), LH (lateral hypothalamus)

**Figure S1.**
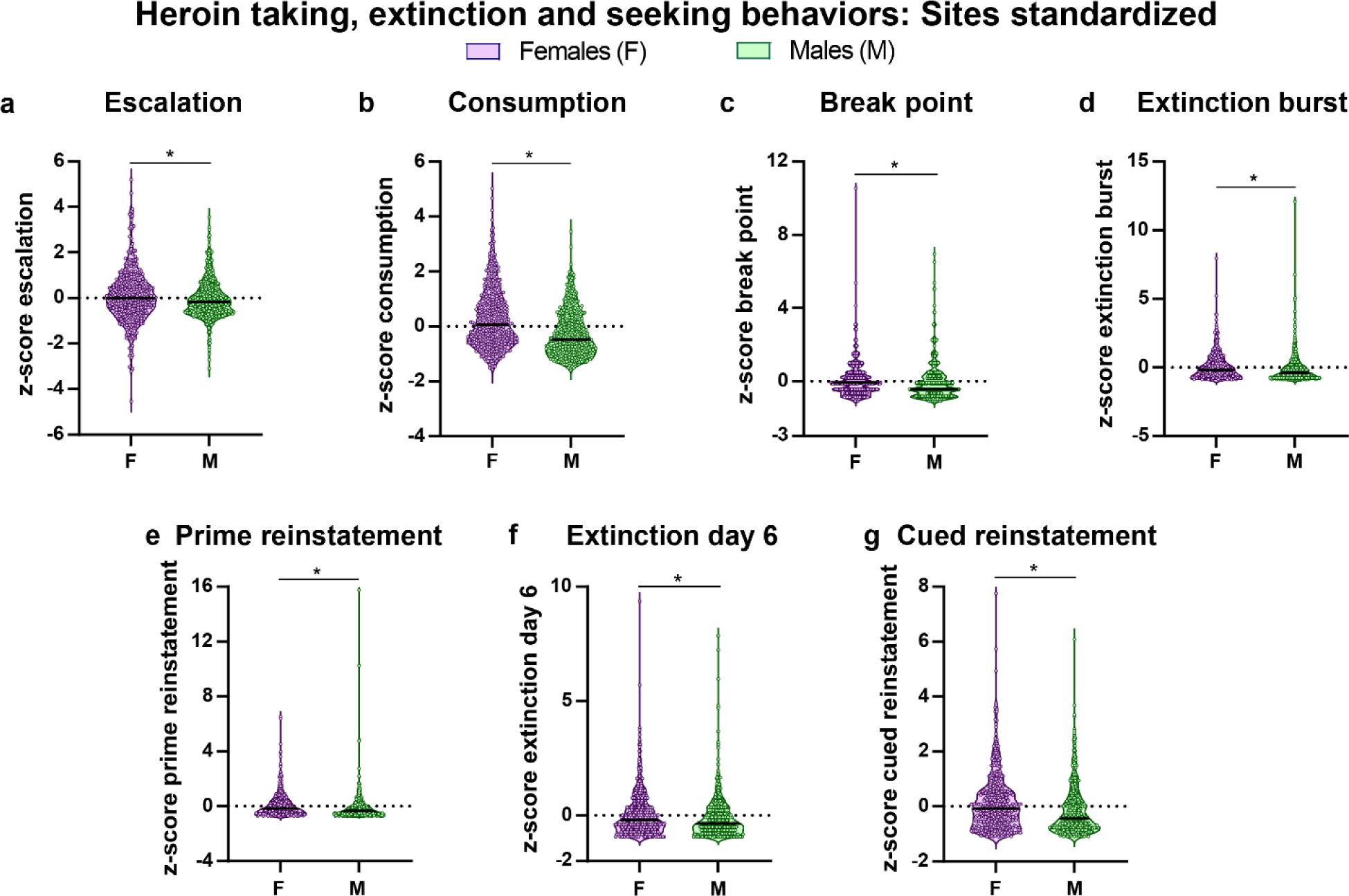
Data for heroin-taking, extinction and seeking behaviors following z-score transformation to standardize data between sites (MUSC vs UCAM). The black bar within each violin plot indicates the median value of the data set, and all data represented as z-scores. Females exhibited potentiated behavioral responding relative to males for **a)** escalation (Mann-Whitney U= 97160, p<0.05) and **b)** consumption (Mann-Whitney U= 71799, p<0.0001) of heroin, **c)** break point achieved during the progressive ratio test (Mann-Whitney U= 92258, p<0.001), extinction behavior during the **d)** first (Mann-Whitney U= 86294, p<0.0001) and **f)** last (Mann-Whitney U= 92872, p<0.002) extinction training, as well as seeking behavior during the **e)** heroin-primed (Mann-Whitney U= 79385, p<0.0001) and **g)** cue-induced (Mann-Whitney U= 85034, p<0.0001) reinstatement. *p<0.05 (Females, n= 452; Males, n=465)

**Figure S2.**
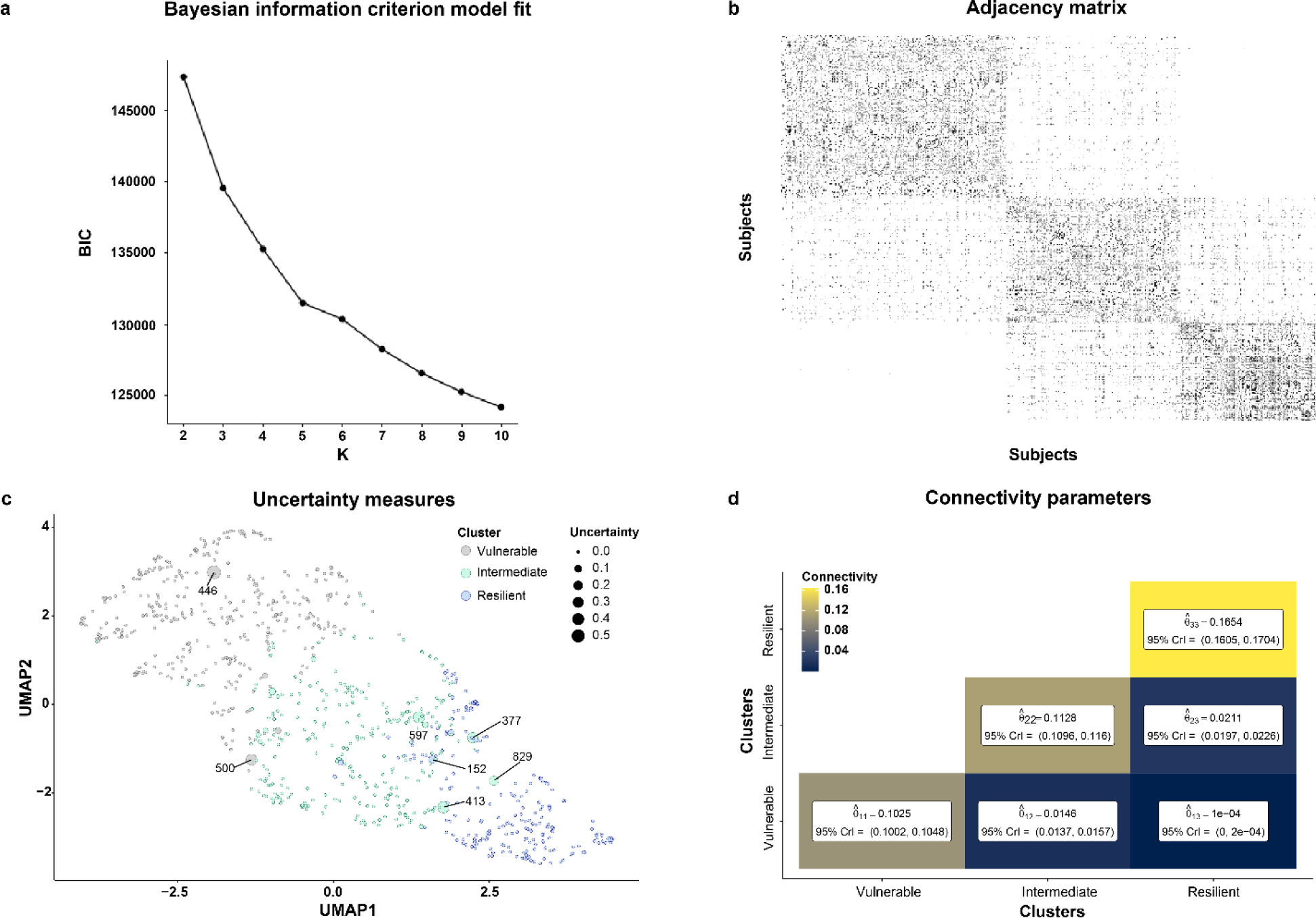
Stochastic block model (SBM) and behavioral clustering parameters. **a)** Bayesian information criterion (BIC) for SBM fits with a range of cluster (K) number between 2 to 10. **b)** Adjacency matrix of inferred clusters from the SBM using K=3 clusters. **c)** Cluster uncertainty score for each rat on UMAP space. Numbers represent rat ID for any rats with less than a 90% certainty of belonging to the assigned cluster (n=7; 0.008% of total population). **d)** Cluster connectivity (ϴ) parameters for within (ϴ_1,1_, ϴ_2,2_, and ϴ_3,3_) and between (ϴ_1,2_, ϴ_1,3_, and ϴ_2,3_) assigned clusters. Higher ϴ value indicates greater connectivity and 95% confidence interval reported below ϴ. Greater connectivity was observed within a cluster relative to between clusters, indicating strong behavioral cohesiveness within SBM cluster analysis assignments. (Vulnerable, n=386; Intermediate, n=300; Resilient, n=231)

**Figure S3.**
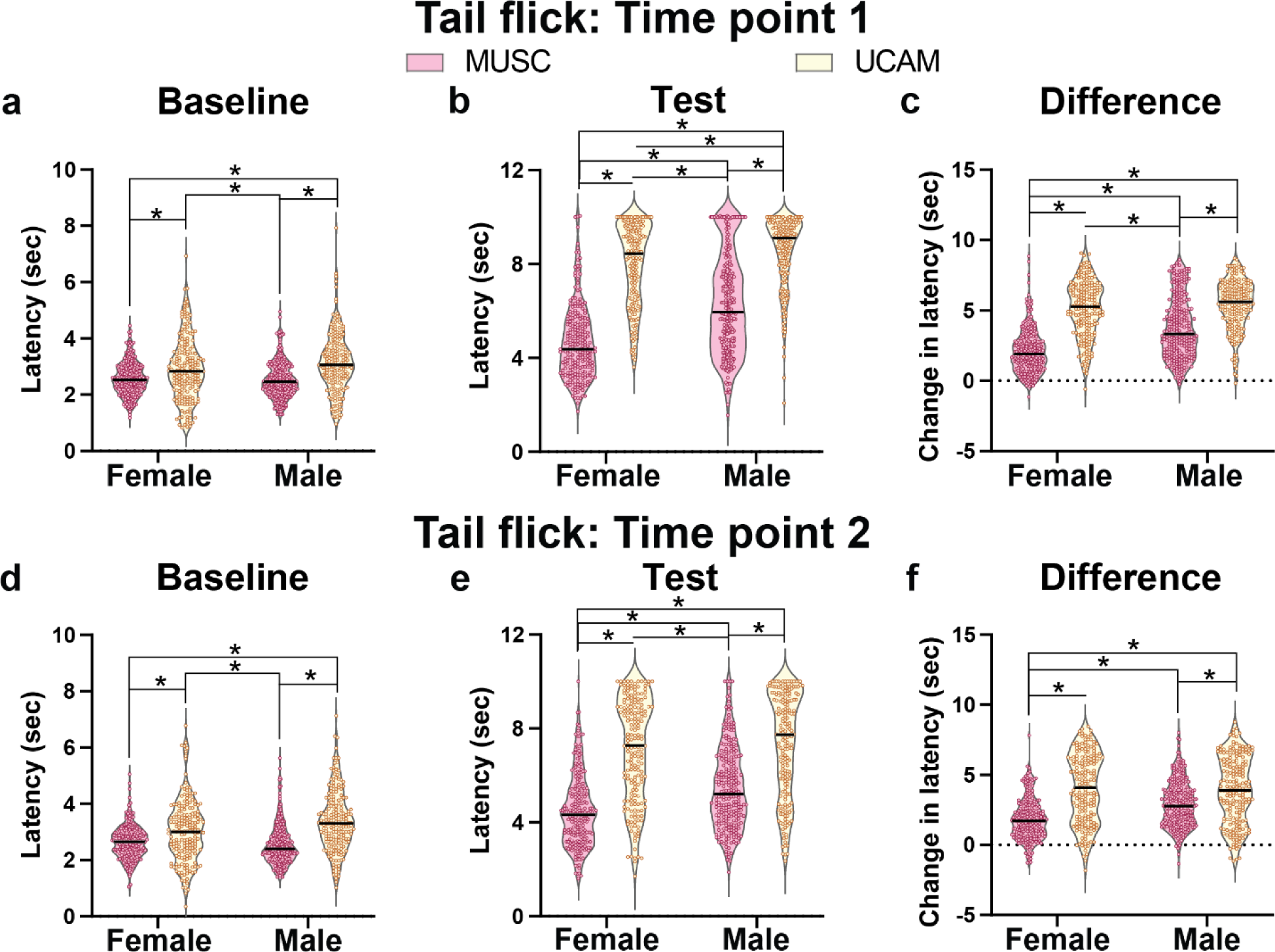
Raw data for tail flick (TF) test behavior for site (MUSC and UCAM) and sex (female and male) in rats. The black bar within each violin plot indicates the median value of the data set. Significant interactions between site and sex were present for both baseline and test sessions prior to (Baseline, **a**, F(1,878)=5.01, p=0.03; Test, **b**, F(1,878)=20.74, p<0.0001; Difference, **c**, F(1,878)=27.63, p<0.0001) and following (Baseline, **a**, F(1,878)=14.36, p=0.0002; Test, **b**, F(1,878)=9.29, p=0.002; Difference, **c**, F(1,878)=21.02, p<0.0001) heroin self-administration training. *p<0.05 (MUSC: Male, n= 253; Female, n= 241; UCAM: Male, n= 194; Female, n=194)

**Figure S4.**
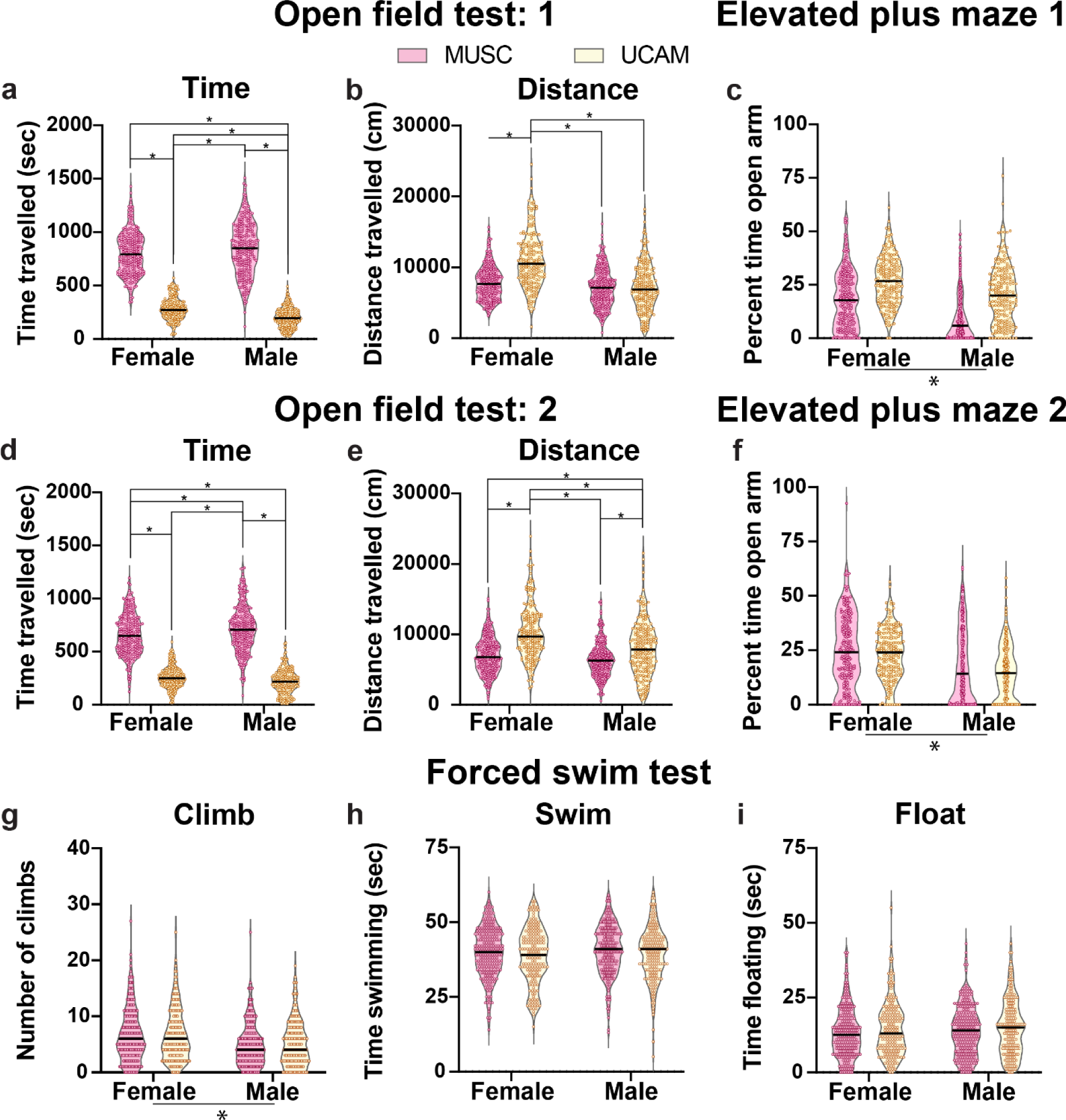
Raw data for open field test (OFT), elevated pluz maze (EPM) and forced swim (FS) test behaviors for site (MUSC and UCAM) and sex (female and male) in rats. Median value for data sets represented as the black bar within each violin plot. A significant testing location and sex interaction was present for time spent moving and distance travelled during the OFT both prior to (Time, **a**, F(1,874)=26.40, p<0.0001; Distance, **b**, F(1,874)=56.95, p<0.0001) and following (Time, **d**, F(1,872)=16.79, p<0.0001; Distance, **e**, F(1,872)=15.67, p<0.0001) heroin self-administration experience. Effects of site and sex, but no interactions, were present for time spent in the open arm of the EPM **c)** before (Site, F(1,877)=126, p<0.0001; Sex, F(1,877)=83.14, p<0.0001) and **f)** after (Site, F(1,877)=4.03, p=0.04; Sex, F(1,877)=52.68, p<0.0001) heroin experience. During the FS test, females exhibited more **g)** climbing behavior relative to males (F(1,843)=26.66, p<0.0001), and rats that were trained at UCAM showed more **h)** swimming behavior (F(1,843)=4.56, p=0.04) compared to those at MUSC. No differences were present for **i)** time spent floating, or interactions for any FST measures. *p<0.05 (n for each test designated as ‘time point 1/time point 2’; OFT: MUSC: Male, n= 253/253; Female, n= 240/241; UCAM: Male, n= 193/189; Female, n=192/193; EPM: MUSC: Male, n= 252/252; Female, n= 241/241; UCAM: Male, n= 194/194; Female, n=194/194; FST: Male, n= 245; Female, n= 236; UCAM: Male, n= 183; Female, n= 182)

**Figure S5.**
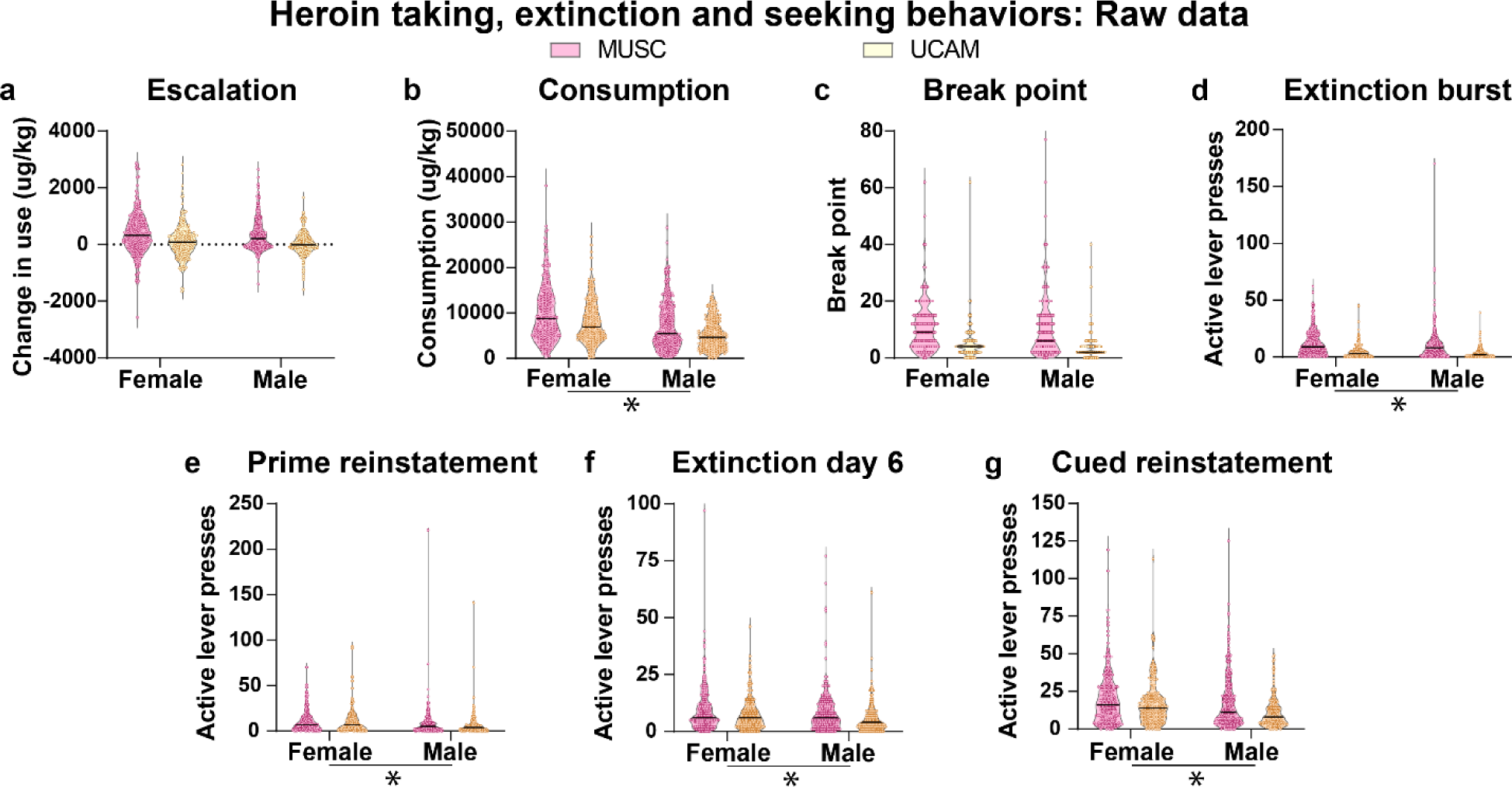
Data for heroin-taking, extinction and seeking behavior for site (MUSC and UCAM) and sex (female and male) in rats. The black bar within each violin plot indicates the median value of the data set, and all data represented as z-scores. Rats undergoing training at MUSC versus UCAM differed in **a)** escalation (F(1,913)=52.17, p<0.0001) and **b)** consumption of heroin (F(1,913)=29.35, p<0.0001), in addition to **c)** break point achieved during the progressive ratio test (F(1,913)=118.40, p<0.0001). Active lever presses made during the **d)** first (F(1,913)=114.40, p<0.0001) and **f)** last day of extinction training (F(1,913)=17.13, p<0.0001), as well as during **g)** cued reinstatement (F(1,913)=21.18, p<0.0001). Relative to males, females exhibited potentiated levels of **b)** heroin consumption (F(1,913)=66.53, p<0.0001), and greater levels of extinction (first day, **d**, F(1,913)=5.46, p=0.02; last day, f, F(1,913)=6.09, p=0.01) and seeking behavior (prime reinstatement, **e**, F(1,913)=15.69, p<0.0001; cued reinstatement, **g**, F(1,913)=15.89, p<0.0001). No significant interactions were present. *p<0.05 (MUSC: Male, n= 271; Female, n= 258; UCAM: Male, n= 194; Female, n=194)

**Figure S6.**
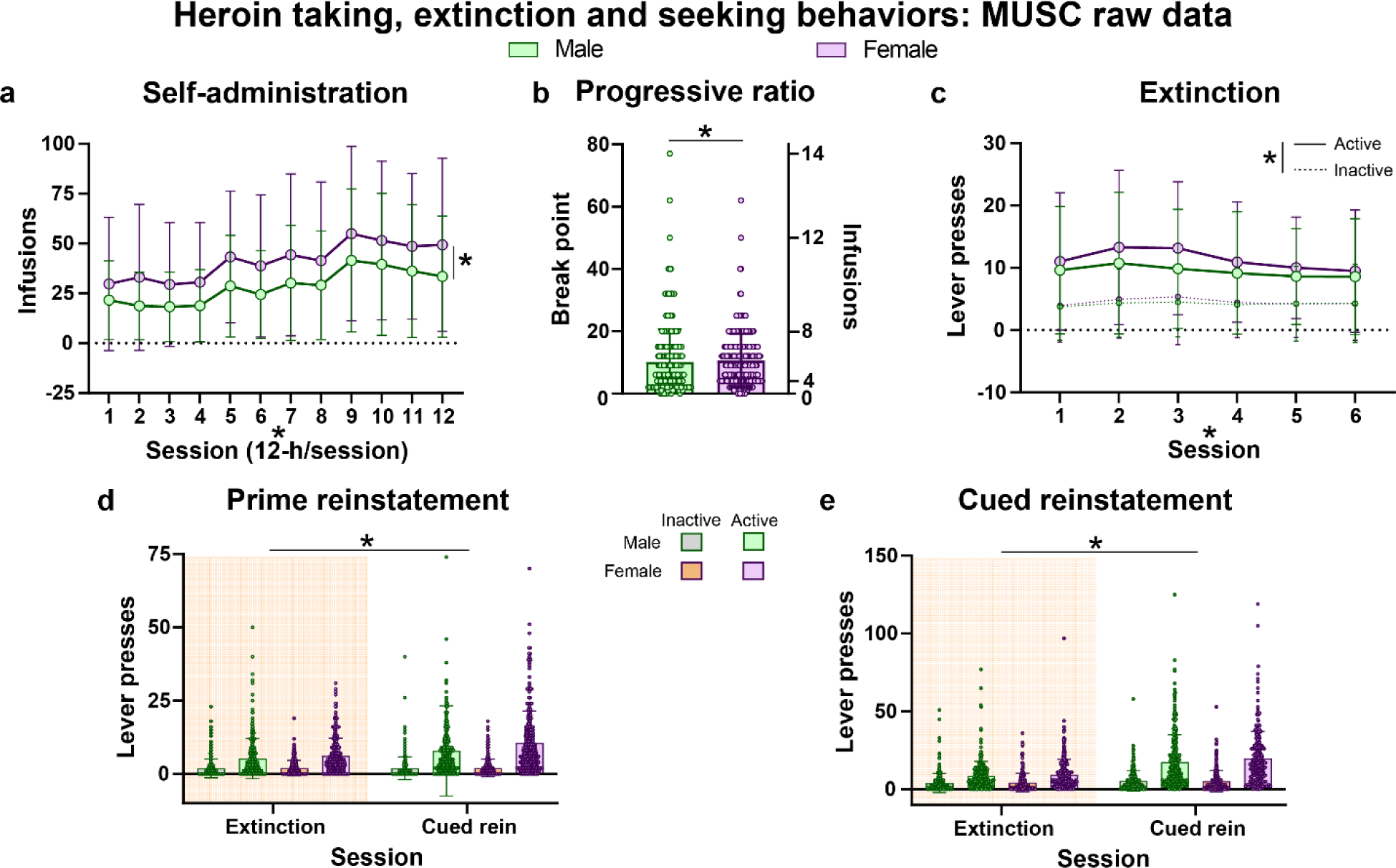
Raw data for heroin-taking, extinction and seeking behavior for male and female rats at MUSC. All data shown with mean ± SD. **a)** All rats escalated intake (20 µg/kg/100 µl infusion over 3 seconds) across heroin self-administration training sessions (2-way repeated ANOVA for session and sex; session, F(5.41,2829)=71.06, p<0.0001), with females maintaining higher levels of consumption relative to males (sex, F(1,525)=39.83, p<0.0001) infusions earned across heroin self-administration training. **b)** Female rats exhibited higher break points during the progressive ratio test compared to males (Mann-Whitney U= 31196, p<0.03). **c)** All rats showed changes in responding on levers during training (3-way repeated ANOVA for session, sex and lever; session x lever, F(5,5245)=7.58, p<0.0001), however, female rats exhibited higher levels of responding on both levers (sex x lever, F(1,1050)=4.03, p<0.05), and across training session (session x sex, F(5,5245)=2.96, p=0.01) compared to males. All rats showed greater responding during the reinstatement test relative to the preceding extinction training session (3-way repeated measures for session, sex and lever;), and differentiated between the levers during both sessions (**d**, lever x session, F(1,1054)=32.95, p<0.0001, all post-hocs p<0.05; **e**, lever x session, F(1,1054)=152.6, p<0.0001, all post-hocs p<0.001). Females showed higher levels of responding during both extinction and prime reinstatement (**d**, sex, F(1,1054)=6.76, p=0.01). The heroin priming injection was 0.25 mg/kg (s.c) given immediately preceding the session. *p<0.05 (Male, n= 271; Female, n= 258)

**Figure S7.**
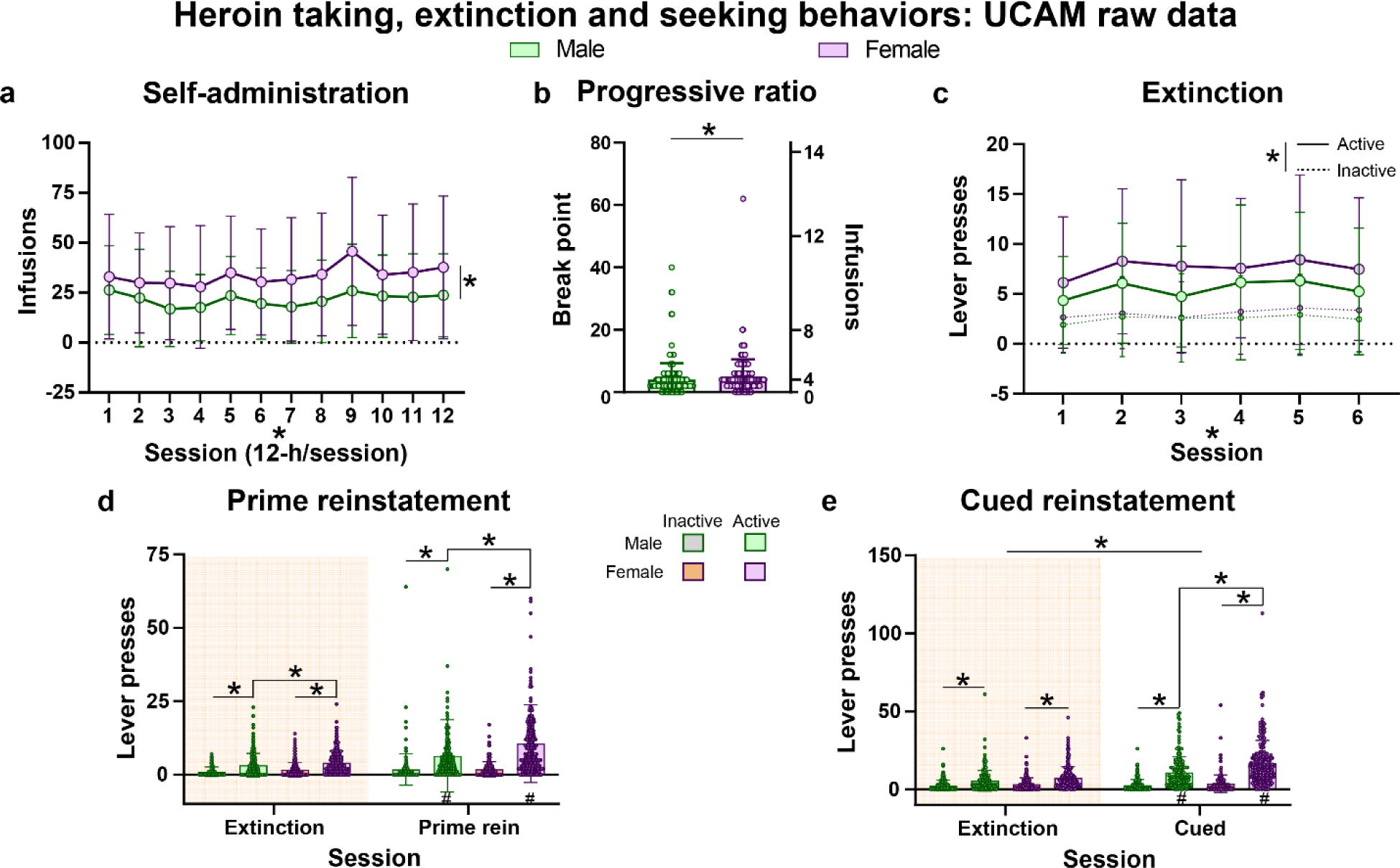
Raw data for heroin-taking, extinction and seeking behavior for male and female rats at UCAM. All data shown with mean ± SD. **a)** Females exhibited higher levels of heroin consumption (20 µg/kg/100 µl infusion over 3 seconds) across all training sessions compared to males (2-way repeated ANOVA for session and sex; session x sex, F(11,4282)=2.53, p=0.04, all post-hoc p<0.02). **b)** Females achieved higher break points during the progressive ratio test compared to male rats (Mann-Whitney U= 15796, p<0.05). **c)** Rats continued to differentiate levers across training (3-way repeated ANOVA for session, sex and lever; session x lever, F(5,3965)=7.58, p=0.03), with females showing higher levels of responding on both levers (sex x lever, F(1,802)=8.01, p=0.005). A lever x session x sex interaction (3-way repeated ANOVA for session, sex and lever) was present for behavior during the extinction session preceding reinstatement and reinstatement for both **d)** prime (F(1,772)=9.54, p<0.02, all post-hocs p<0.05) and **e)** cue (F(1,772)=9.60, p=0.002, all post-hocs p<0.005). The heroin priming injection was 0.25 mg/kg (s.c) given immediately preceding the session. *p<0.05 (Male, n= 194; Female, n= 194)

**Figure S8.**
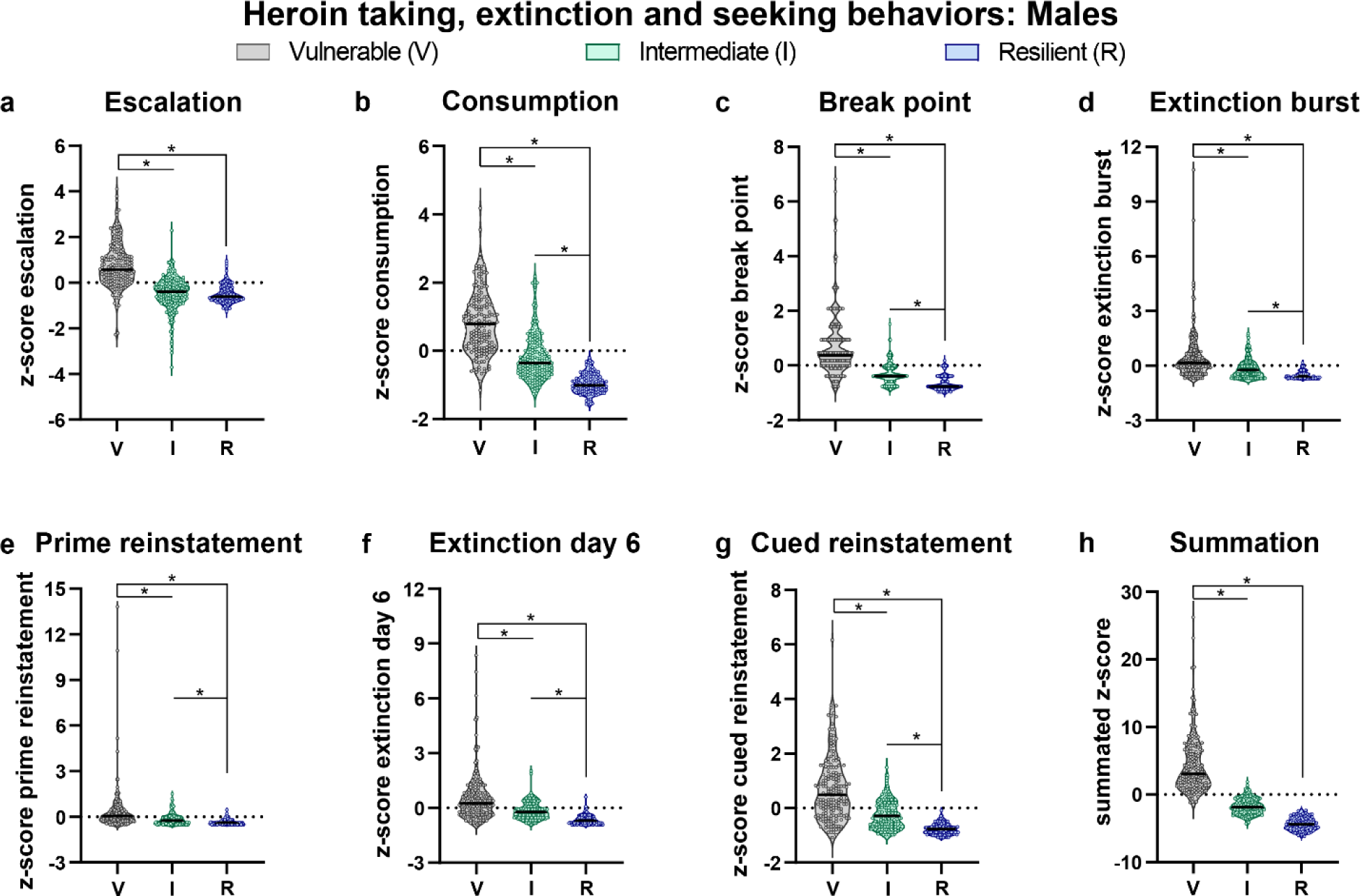
Data for heroin-taking, extinction and seeking behavior for OUD vulnerability clusters in male rats. The black bar within each violin plot indicates the median value of the data set, and all data represented as z-scores. Behavior differed across clusters for **a)** escalation (Kruskal-Wallis, H(2)=206.9, p<0.0001) and **b)** consumption (Kruskal-Wallis, H(2)=278.2, p<0.0001) of heroin, **c)** break point achieved during the progressive ratio test (Kruskal-Wallis, H(2)=240.5 p<0.0001), extinction behavior during the **d)** first (Kruskal-Wallis, H(2)=140.7, p<0.0001) and **f)** last (Kruskal-Wallis, H(2)=177.7, p<0.0001) extinction training session and seeking behavior during the **e)** heroin-primed (Kruskal-Wallis, H(2)=115.7, p<0.0001) and **g)** cued reinstatement test (Kruskal-Wallis, H(2)=212.6, p<0.0001). **h)** summation of z-scores for the behaviors in a-g (Kruskal-Wallis, H(2)=388.6, p<0.0001). All significant post-hoc tests were p<0.0001. *p<0.05 (Vulnerable, n= 182; Intermediate, n= 168; Resilient, n=115)

**Figure S9.**
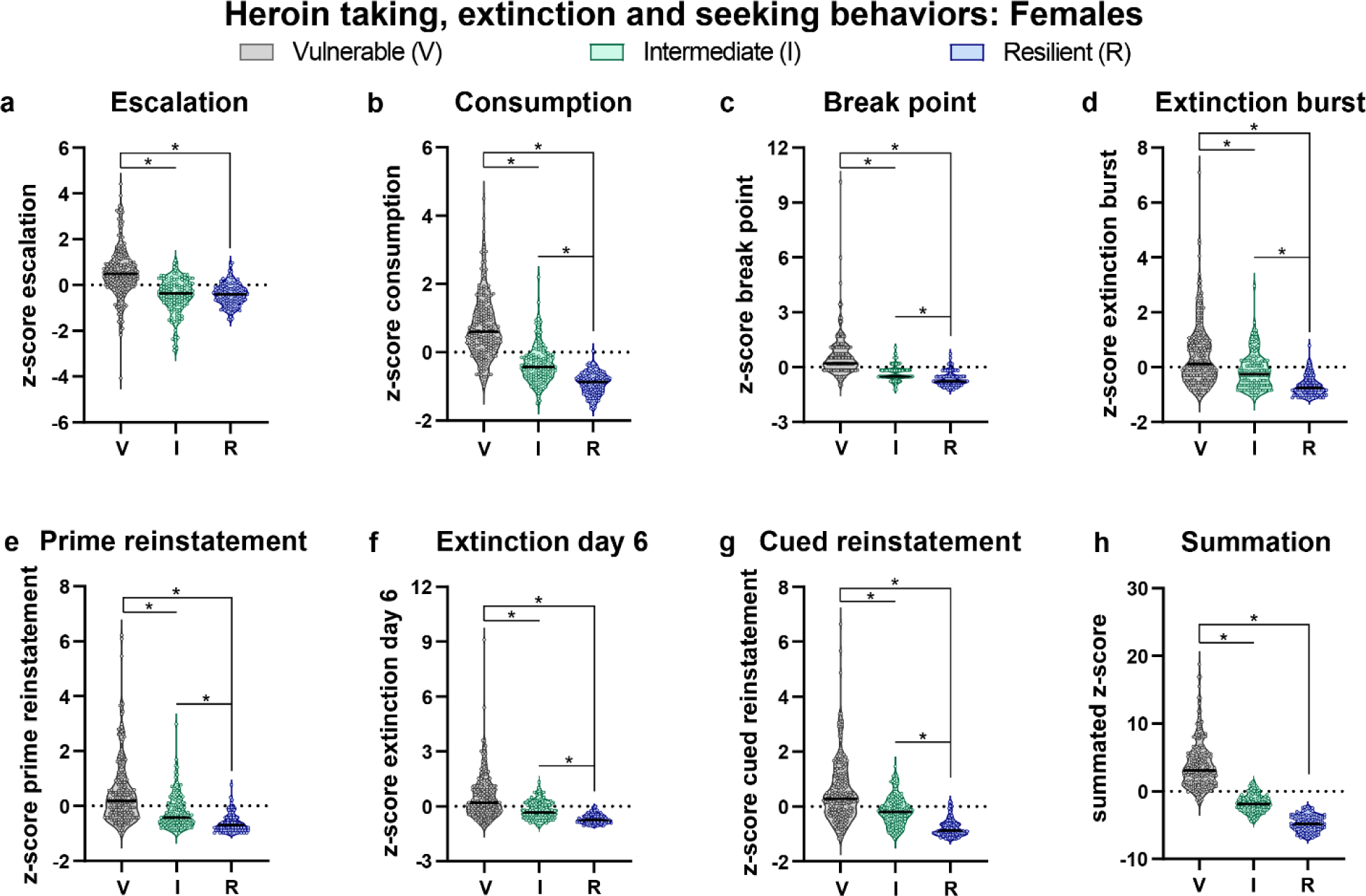
Data for heroin-taking, extinction and seeking behavior for OUD vulnerability clusters in female rats. The black bar within each violin plot indicates the median value of the data set, and all data represented as z-scores. Behavioral differences are present in clusters for **a)** escalation (Kruskal-Wallis, H(2)=133.9, p<0.0001) and **b)** consumption (Kruskal-Wallis, H(2)=274.8, p<0.0001) of heroin, **c)** break point achieved during the progressive ratio test (Kruskal-Wallis, H(2)=230.4 p<0.0001), extinction behavior during the **d)** first (Kruskal-Wallis, H(2)=135.5, p<0.0001) and **f)** last (Kruskal-Wallis, H(2)=171.9, p<0.0001) extinction training session and seeking behavior during the **e)** heroin-primed (Kruskal-Wallis, H(2)=150.0, p<0.0001) and **g)** cued reinstatement test (Kruskal-Wallis, H(2)=191.8, p<0.0001). **h)** summation of z-scores for the behaviors in a-g (Kruskal-Wallis, H(2)=372, p<0.0001). All significant post-hoc tests were p<0.0001. *p<0.05 (Vulnerable, n= 204; Intermediate, n= 132; Resilient, n=116)

**Figure S10.**
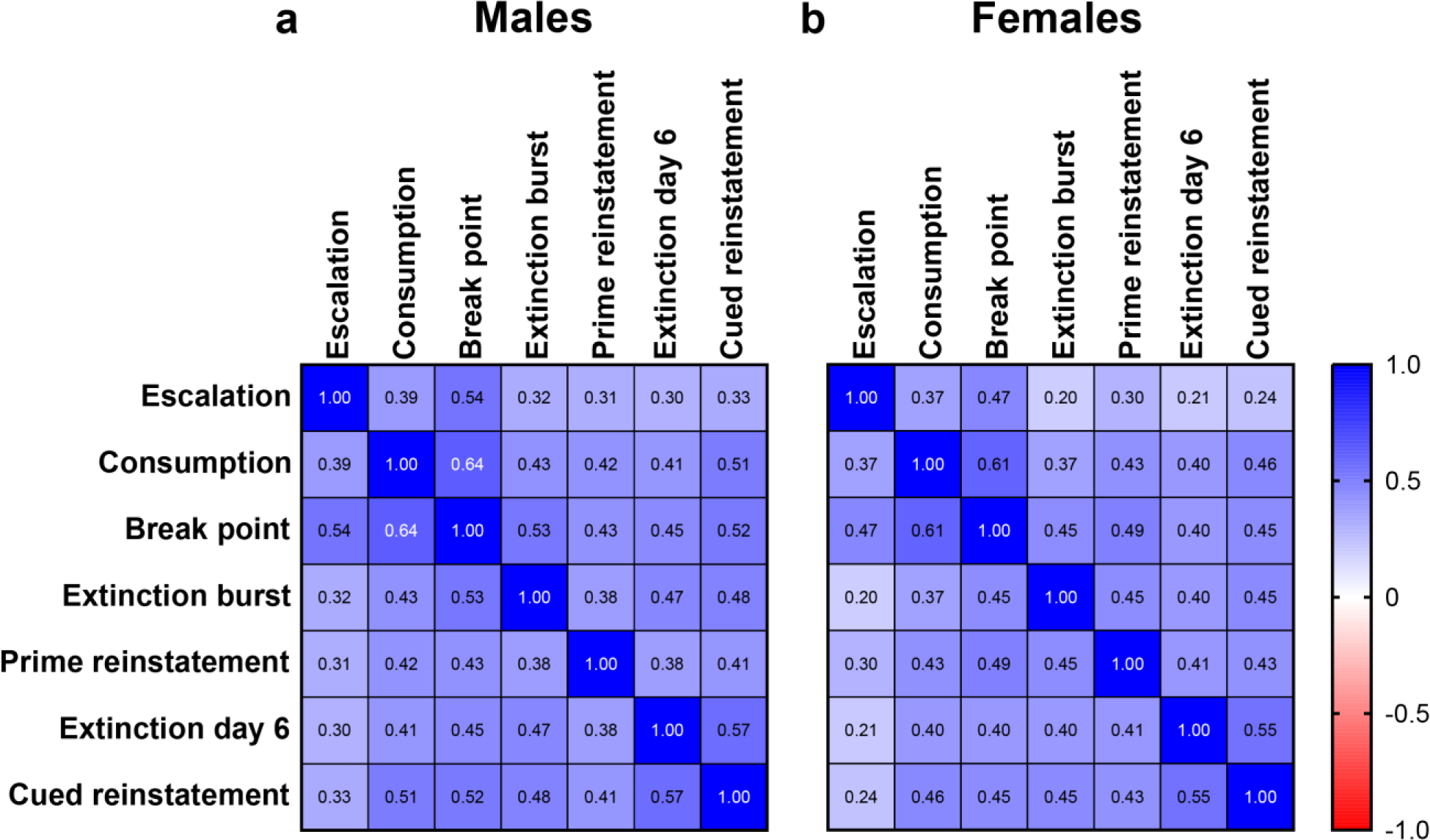
Correlation matrixes for behaviors used in non-linear clustering analysis for male and female rats. The value within each box is the Spearman’s correlation coefficient. Significant covariance was present for all behavioral interactions for both **a)** male and **b)** female rats. p<0.05 (Male, n= 465; Female, n= 452)

**Figure S11.**
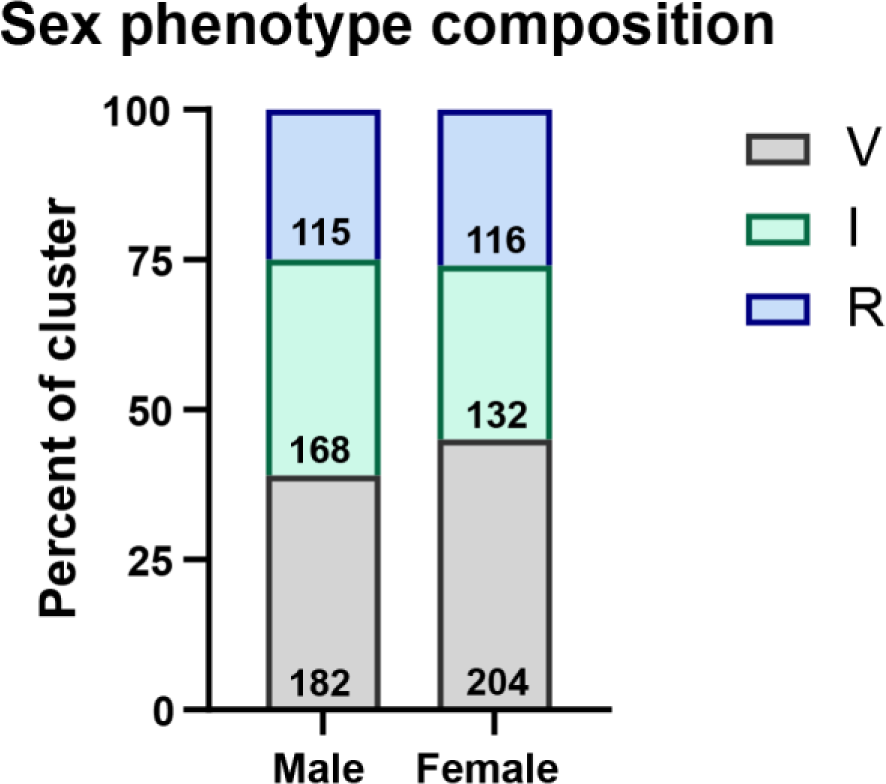
OUD phenotypic composition for male and female vulnerable (V), intermediate (I) and resilient (R) rats. Cluster composition within sex did not differ between male and female rats (x^2^(1)=5.40, p=0.07). Number inside of bar is the number of animals for that cluster.

**Figure S12.**
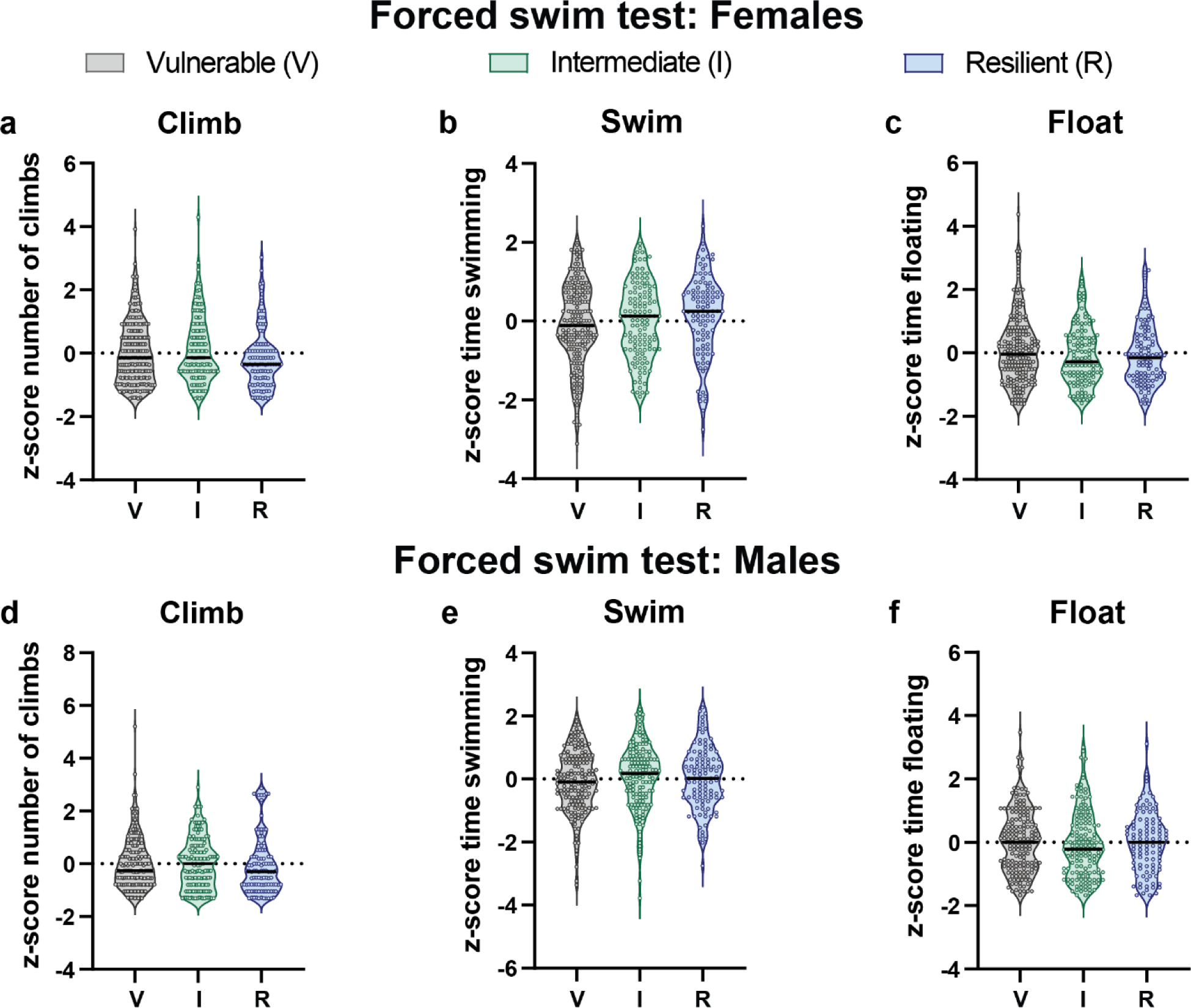
Forced-swim (FS) test behavior for OUD vulnerability clusters in female and male rats. Rats underwent a FS test prior to heroin experience. The black bar within each violin plot indicates the median value of the data set, and all data represented as z-scores. Clusters did not differ from one another for either female or male rats in **a, d)** number of climbs (Females: Kruskal-Wallis, H(2)=4.1, p=0.13; Males: Kruskal-Wallis, H(2)=3.3, p=0.19), **b, e)** time spent swimming (Females: Kruskal-Wallis, H(2)=3.5, p=0.18; Males: Kruskal-Wallis, H(2)=2.1, p=0.35), **c, f)** or time spent floating (Females: Kruskal-Wallis, H(2)=1.7, p=0.43; Males: Kruskal-Wallis, H(2)=2.1, p=0.36) for either females or males. (Females, n= 418; Males, n=429)

**Figure S13.**
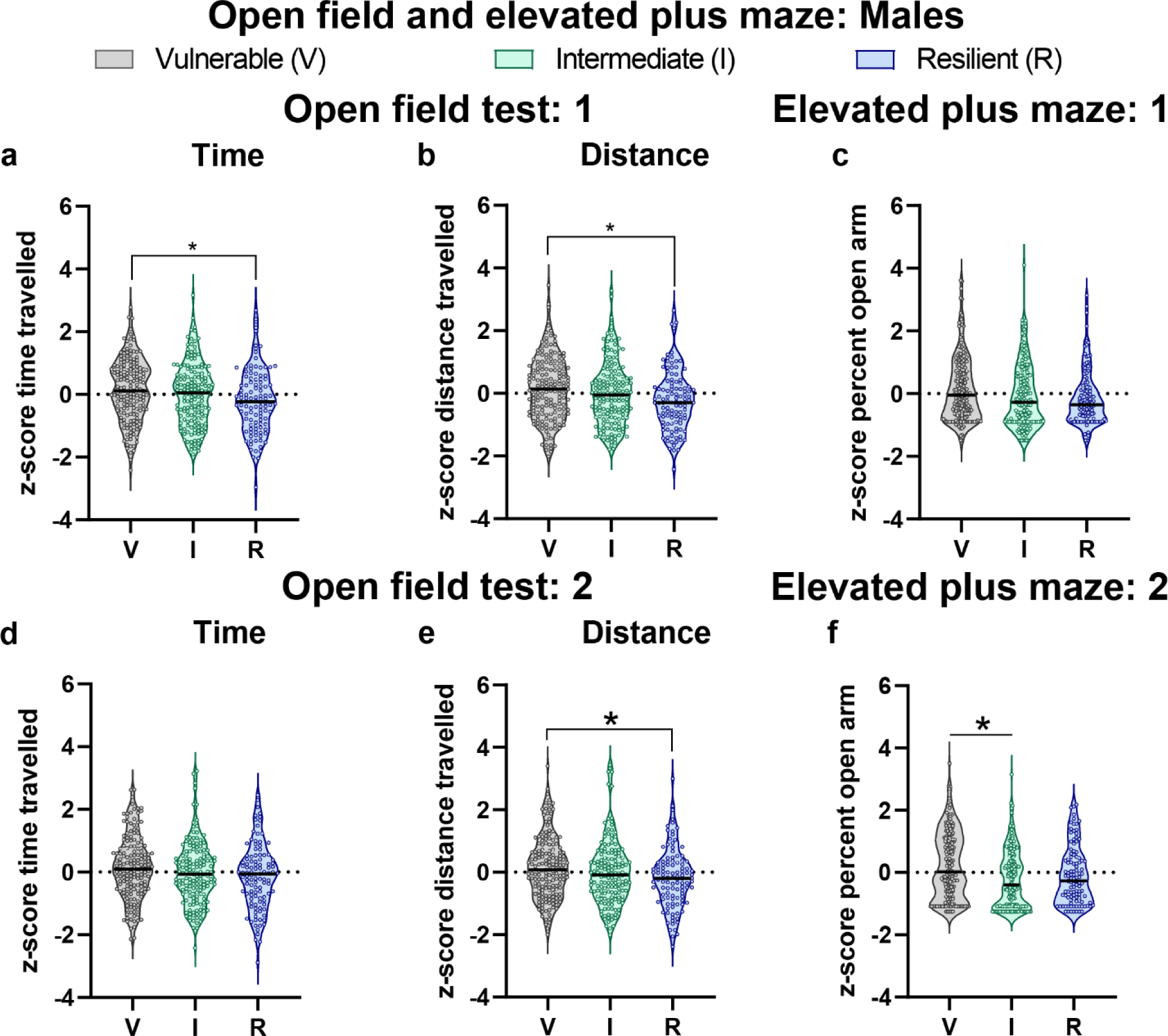
Open field test and elevated plus maze behavior for OUD vulnerability clusters in male rats. The black bar within each violin plot indicates the median value of the data set, and all data represented as z-scores. Vulnerable rats spent more **a)** time moving (ANOVA, F(2,442)=4.19, p=0.02; post-hoc p=0.01) and **b)** travelled a greater distance (ANOVA, F(2,442)=5.34, p=0.001; post-hoc p=0.01) than resilient rats during the open field task prior to heroin experience. Following heroin experience, rats did not differ in **d)** time spent moving (ANOVA, F(2,439)=2.70, p=0.07), but vulnerable rats maintained **e)** higher levels of distance travelled relative to resilient rats (Kruskal-Wallis, H(2)=7.5, p=0.02; post-hoc p=0.02). Phenotypes overall differed in percent time spent in the open arm of the elevated plus maze **c)** prior to (Kruskal-Wallis, H(2)=6.6, p=0.04, no significant post-hocs) and **f)** following (Kruskal-Wallis, H(2)=14.2, p=0.0008; vulnerable vs intermediate post-hoc p=0.0005) heroin experience. *p<0.05 (OFT: Vulnerable, n=175 (174 OFT 2); Intermediate, n=161 (159 OFT 2); Resilient, n=109; EPM: Vulnerable, n= 175 (174 OFT 2); Intermediate, n= 163 (161 OFT 2); Resilient, n=109)

**Figure S14.**
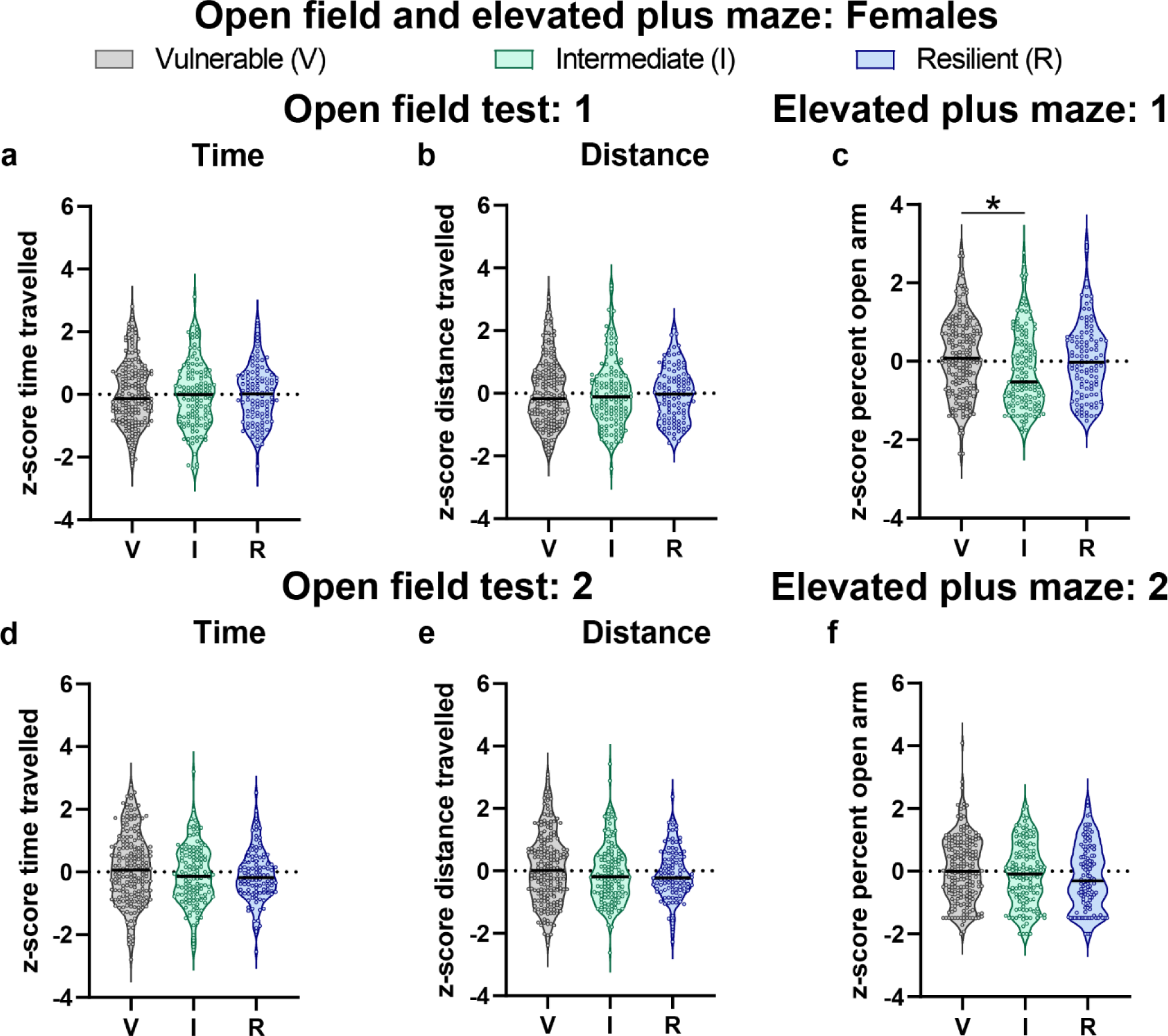
Open field test and elevated plus maze behavior for OUD vulnerability clusters in female rats. The black bar within each violin plot indicates the median value of the data set, and all data represented as z-scores. Rats did not differ in time spent moving or total distance travelled prior to (**a)** Time: Kruskal-Wallis, H(2)=0.21, p=0.90; **b)** Distance: Kruskal-Wallis, H(2)=0.22, p=0.89) and following (**d)** Time: F(2,432)=2.88, p=0.06; **e)** Distance: Kruskal-Wallis, H(2)=3.1, p=0.22) heroin experience. **c)** A difference in behavior between the phenotypes were present in EPM 1 (Kruskal-Wallis, H(2)=11.8, p=0.003), with intermediate rats spending less percent time in the open arm compared to vulnerable rats (p=0.002). **f)** No phenotypic differences were observed in the second EPM (F(2,432)=2.06, p=0.13). *p<0.05 (OFT: Vulnerable, n= 192 (198 OFT 2); Intermediate, n= 130; Resilient, n=107; EPM: Vulnerable, n= 198; Intermediate, n= 130; Resilient, n=107)

**Figure S15.**
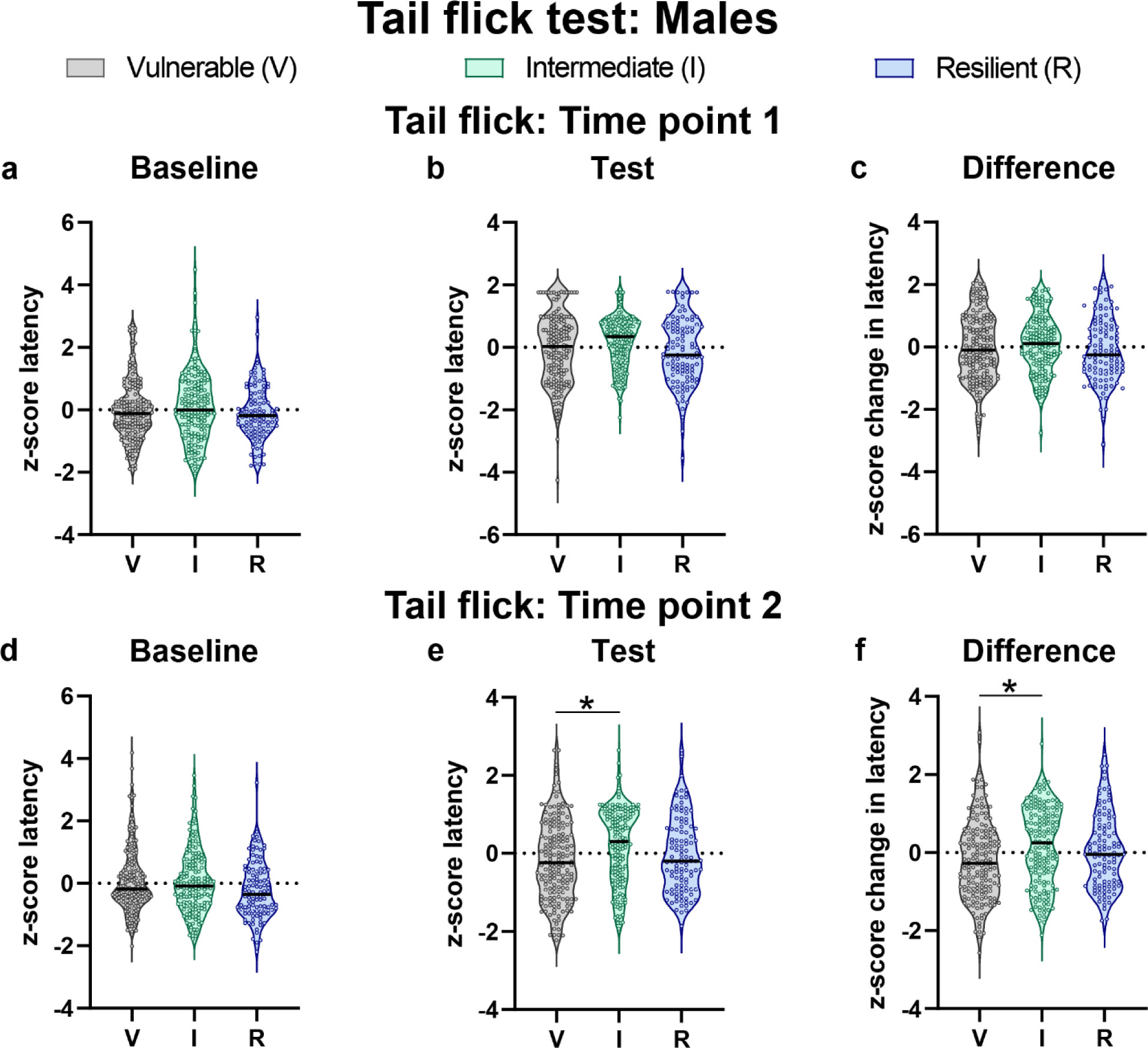
Tail flick test behavior for OUD vulnerability clusters in male rats. The black bar within each violin plot indicates the median value of the data set, and all data represented as z-scores. No phenotypics differences were observed in the **a)** baseline session (Kruskal-Wallis, H(2)=2.8, p=0.24), and though a difference in phenotype was present for the **b)** test session (Kruskal-Wallis, H(2)=6.6, p=0.04), there were no significant post hoc tests. **c)** A difference in latency between the two sessions were not observed (F(2,444)=1.49, p=0.23). Phenotypes did not differ in behavior in the **d)** baseline session following heroin experience (Kruskal-Wallis, H(2)=5.9, p=0.05), but did in the **e)** test session (Kruskal-Wallis, H(2)=13.5, p=0.001) with vulnerable rats having lower latency times than intermediate rats (p=0.001). **f)** This relationship was present in the latency difference between the two sessions (Kruskal-Wallis, H(2)=11.9, p=0.003), with intermediate rats having a higher latency than vulnerable rats (p=0.002). *p<0.05 (Vulnerable, n= 175; Intermediate, n= 163; Resilient, n=109)

**Figure S16.**
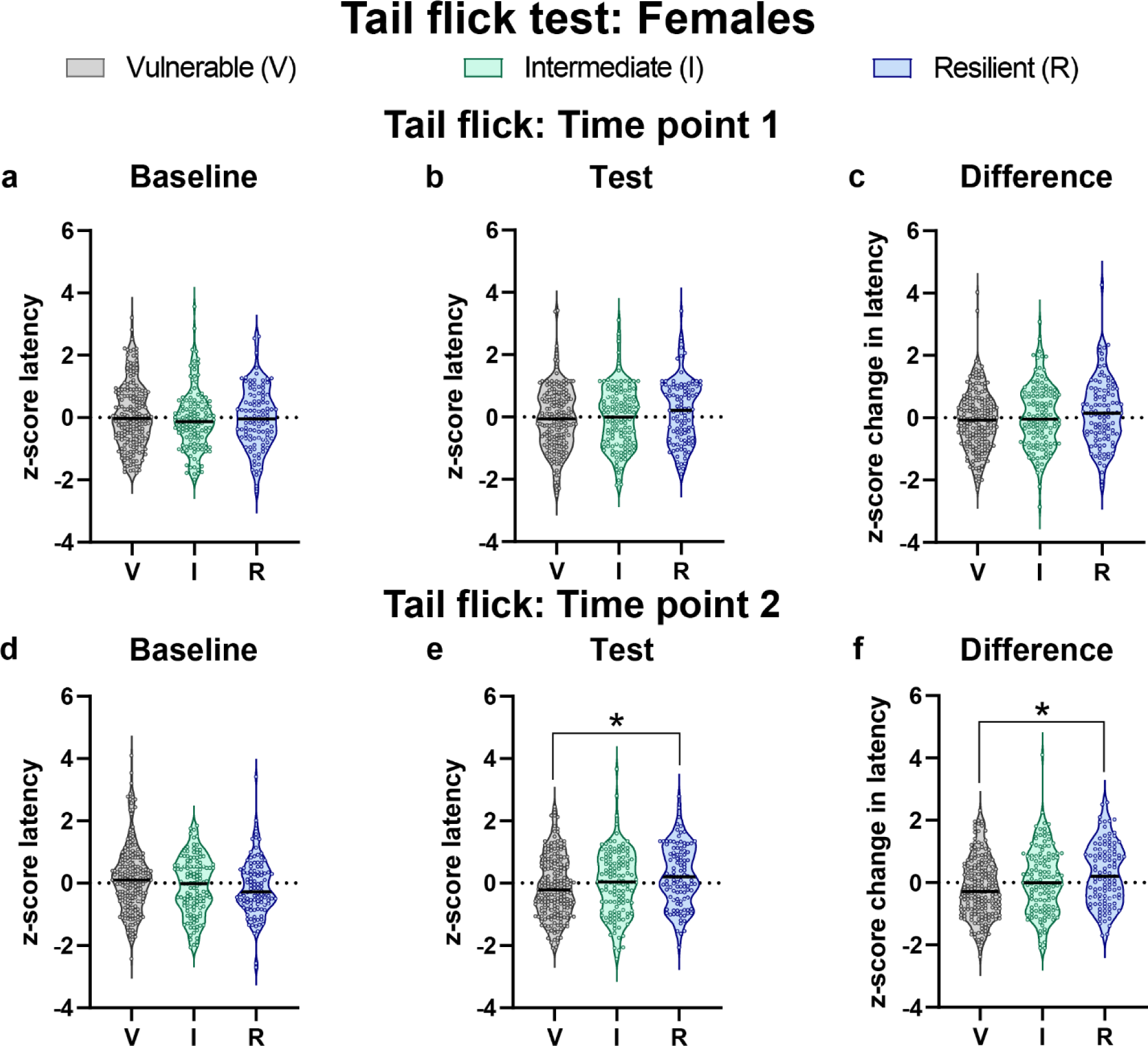
Tail flick test behavior for OUD vulnerability clusters in female rats. The black bar within each violin plot indicates the median value of the data set, and all data represented as z-scores. No phenotypic differences were observed in the **a)** baseline session (Kruskal-Wallis, H(2)=1.6, p=0.46) and **b)** test session (Kruskal-Wallis, H(2)=3.7, p=0.16), or the **c)** latency difference between the two sessions (F(2,432)=2.02, p=0.13) prior to heroin training. Following heroin experience, clusters did not differ during the **d)** baseline session (Kruskal-Wallis, H(2)=5.3, p=0.07), however, vulnerabe rats showed lower latency responses compared to the resilient rats during the **e)** test session (Kruskal-Wallis, H(2)=9.5, p=0.009; vulnerable vs intermdiate p=0.006) and **f)** the latency difference between the two sessions (F(2,432)=7.76, p=0.0005). *p<0.05 (Vulnerable, n= 198; Intermediate, n= 130; Resilient, n=107)

**Figure S17.**
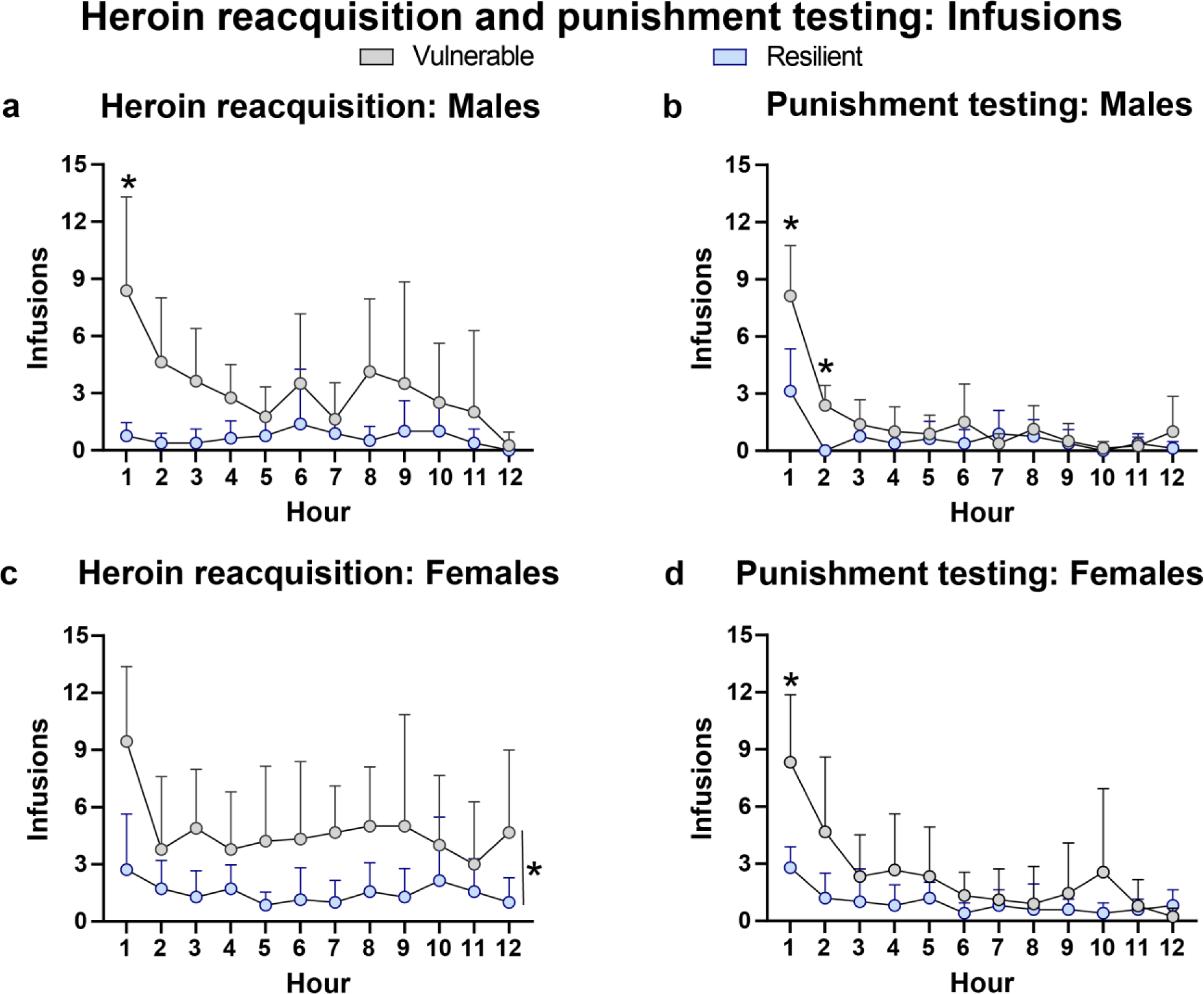
Infusions earned across the last day of heroin reacquisition and the punishment test. Data analyzed with a 2-way repeated ANOVA for phenotype and hour and shown as mean + SD. Vulnerable animals (males, n=8; females, n=7) maintained higher levels of infusions earned across the last day of heroin reacquisition training (session 3) relative to resilient animals (males, n=8; females, n=7), with **a)** effects localized to the first hour of the session in males (hour x cluster, F(11,154)=2.64, p=0.004) and **d)** across training for females (cluster, F(1,14)=12.65, p=0.003). These effects were maintained during the punishement test, with **b)** vulnerable males earning more infusions in the first two hours of the test (hour x cluster, F(11,154)=7.05, p<0.001), and female vulnerable rats in the first hour of the test (hour x cluster, F(11,132)=2.59, p=0.005). *p<0.05

**Figure S18.**
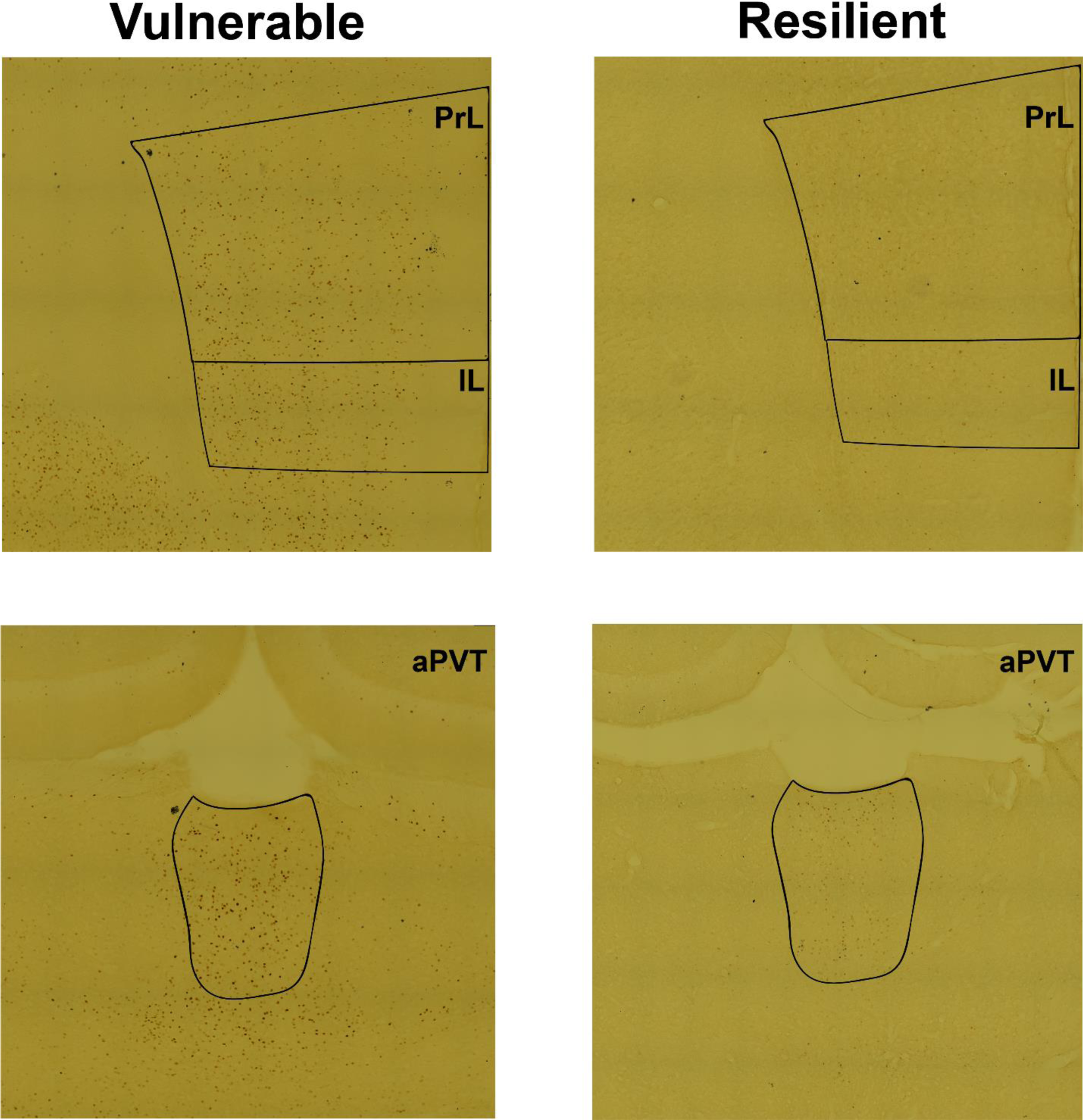
Representative micrographs of cue-induced Fos protein density in vulnerable and resilient rats. Images are from regions that showed phenotypic differences in Fos protein quantification. PrL (prelimbic cortex), IL (infralimbic cortex); aPVT (anterior paraventricular nucleus of the thalamus)

**Figure S19.**
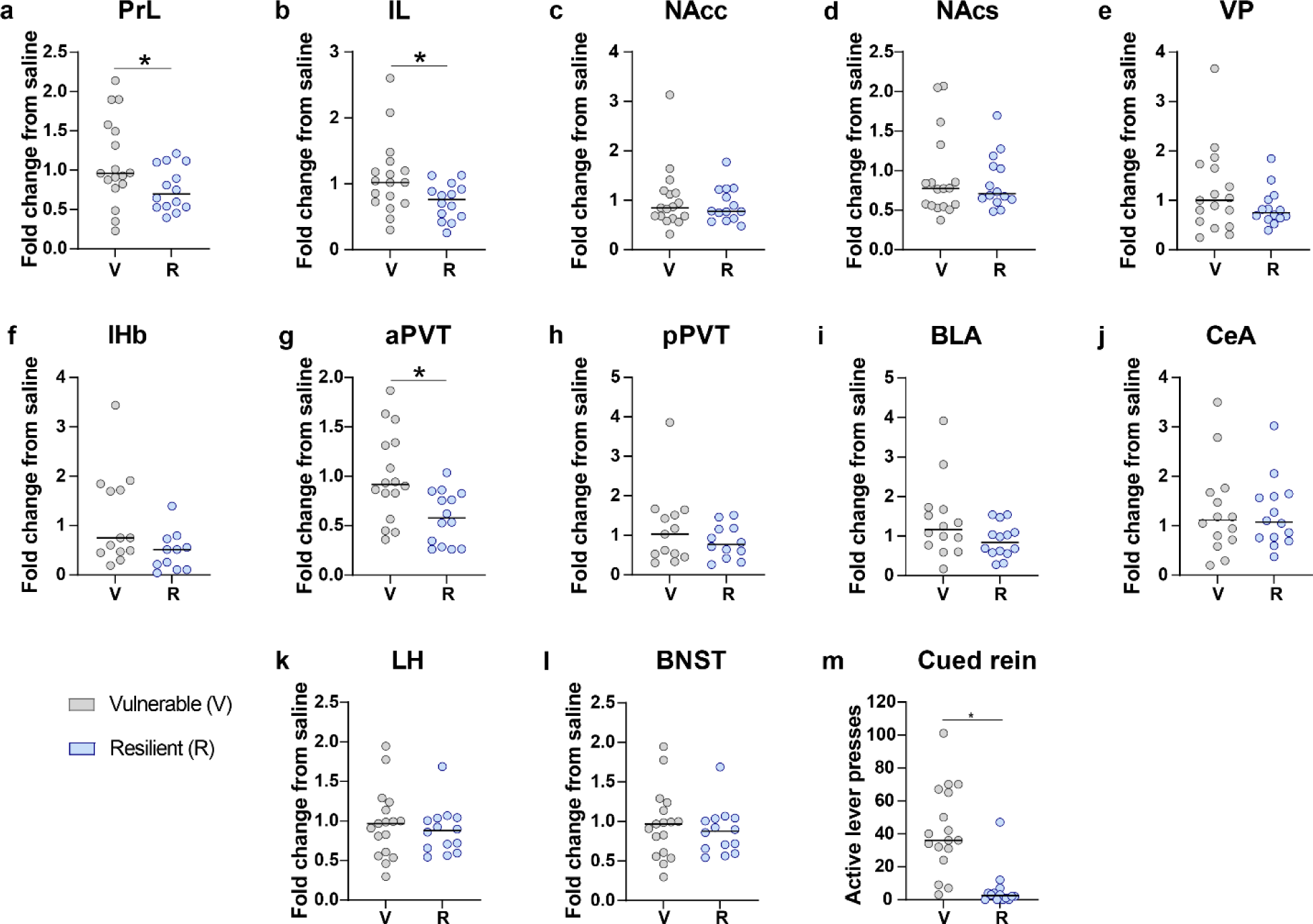
Fos protein quantification comparisons between OUD vulnerable and resilient rats. All data was standardized to saline control rats prior to comparison. Relative to rats in the vulnerable cluster, resilient rats exhibited hypofunctioning in the **a)** PrL, **b)** IL and **g)** aPVT. Phenotypes did not differ in Fos protein quantity in other ROIs. **m)** Vulnerable animals maintained higher levels of cued seeking behavior during the second cued reinstatement test compared to resilient animals (Mann-Whitney U= 18.50, p<0.0001). The anterior-posterior (AP) coordinate from the skull relative to Bregma (mm) where quantification of select ROIs occurred are within the parantheses. PrL (prelimbic cortex, AP 3), IL (infralimbic cortex, AP 3), NAcc (nucleus accumbens core, AP 1.56), NAcs (nucleus accumbens shell, AP 1.56), VP (ventral pallidum, AP 0), lHb (lateral habenula, AP −3), aPVT (anterior paraventricular nucleus of the thalamus, AP −2), pPVT (posterior paraventricular nucleus of the thalamus, AP −3), BLA (basolateral amygdala, AP −2.4), CeA (central amygdala, AP −2.4), LH (lateral hypothalamus, AP −2.4), BNST (bed nucleus of the stria terminalis, AP 0). *p<0.05 (Vulnerable, n= 17; Resilient, n= 14; Saline, n=11)

**Figure S20.**
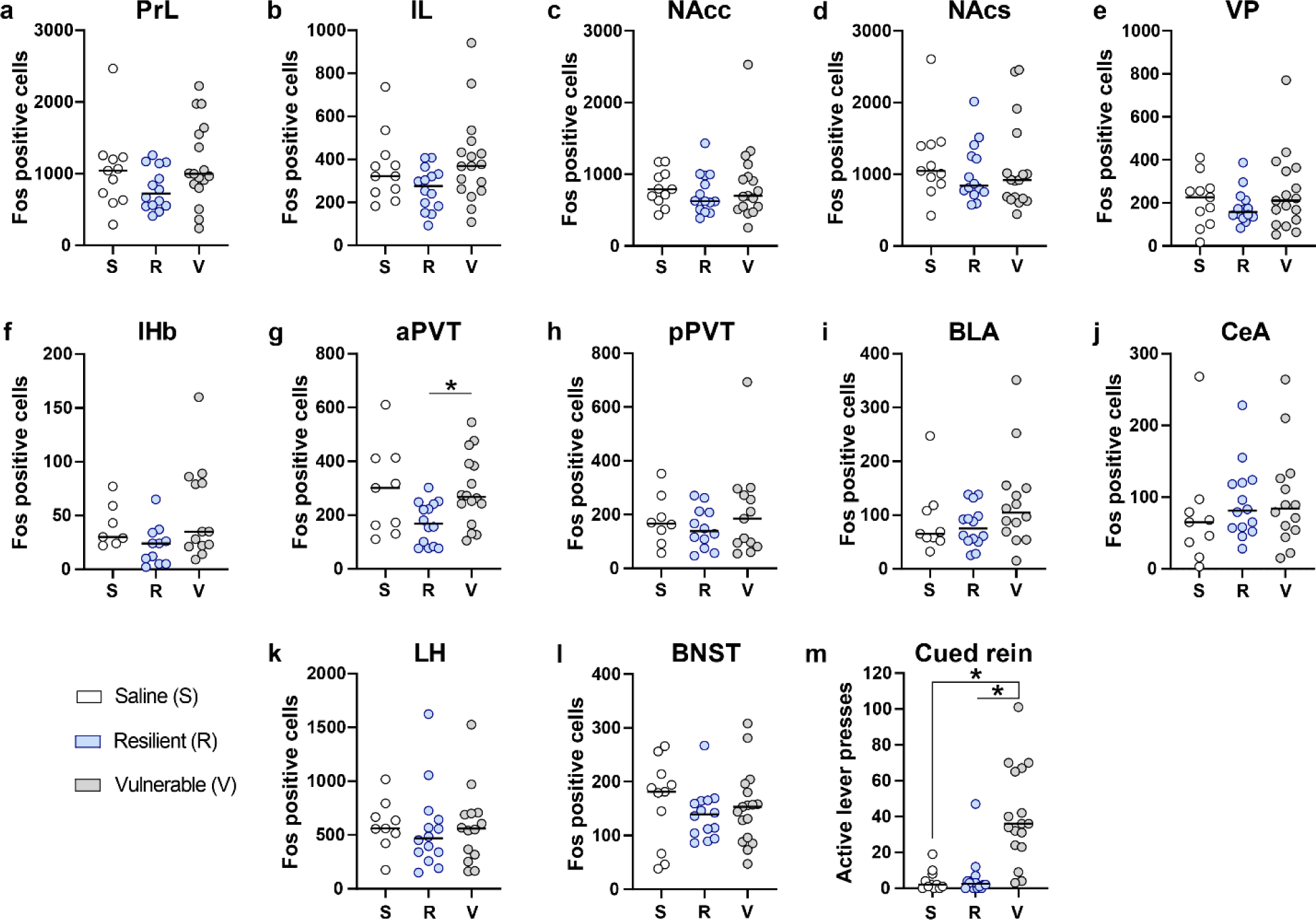
Raw Fos protein cell quantification comparisons between saline control, OUD resilient and vulnerable rats. Relative to rats in the vulnerable group, resilient rats exhibited lower Fos labelling in the **g)** aPVT (F(2,36)= 4.18, p=0.02, post-hoc p=0.03). Groups did not differ in Fos protein quantity in other ROIs. **m)** Vulnerable animals maintained higher levels of cued seeking behavior during the second cued reinstatement test compared to saline control and resilient animals (Kruskal-Wallis, H(2)= 21.96, p<0.0001, post-hocs p=0.0002). The anterior-posterior (AP) coordinate from the skull relative to Bregma (mm) where quantification of select ROIs occurred are within the parantheses. PrL (prelimbic cortex, AP 3), IL (infralimbic cortex, AP 3), NAcc (nucleus accumbens core, AP 1.56), NAcs (nucleus accumbens shell, AP 1.56), VP (ventral pallidum, AP 0), lHb (lateral habenula, AP −3), aPVT (anterior paraventricular nucleus of the thalamus, AP −2), pPVT (posterior paraventricular nucleus of the thalamus, AP −3), BLA (basolateral amygdala, AP −2.4), CeA (central amygdala, AP −2.4), LH (lateral hypothalamus, AP −2.4), BNST (bed nucleus of the stria terminalis, AP 0). *p<0.05 (Vulnerable, n= 17; Resilient, n= 14; Saline, n=11)

